# Neuron-astrocyte metabolic coupling during neuronal stimulation protects against fatty acid toxicity

**DOI:** 10.1101/465237

**Authors:** Maria S. Ioannou, Jesse Jackson, Shu-Hsein Sheu, Chi-Lun Chang, Aubrey V. Weigel, Hui Liu, H. Amalia Pasolli, C. Shan Xu, Song Pang, Harald F. Hess, Jennifer Lippincott-Schwartz, Zhe Liu

## Abstract

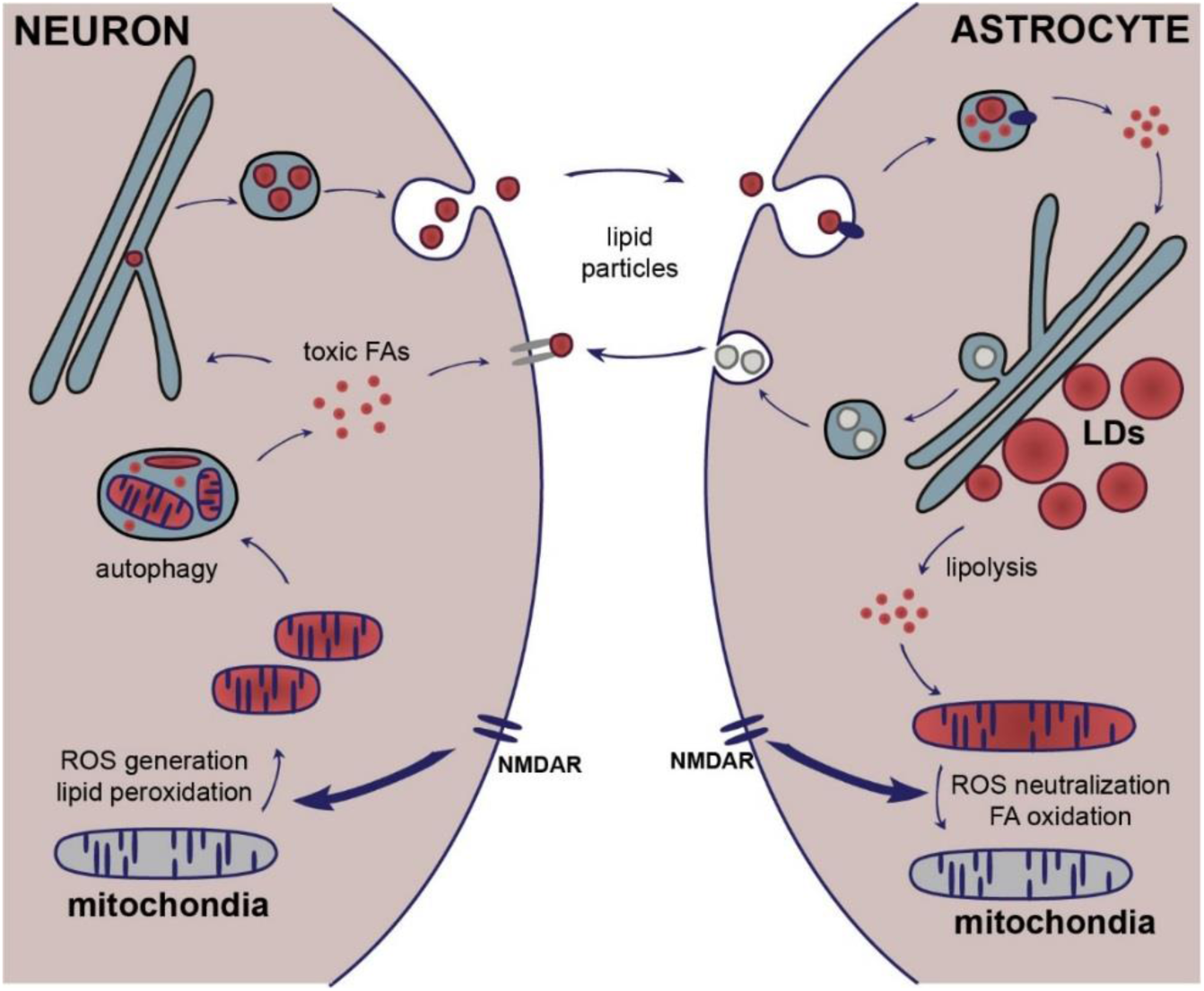

**HIGHLIGHTS:**

- Hyperactive neurons generate and release excess peroxidated fatty acids (FAs)
- Astrocytes endocytose toxic FAs as lipoprotein particles, delivering them to lipid droplets
- Lipid droplets trigger upregulation of genes involved in metabolism and detoxification
- Astrocytes detoxify and consume FAs by mitochondrial oxidation in response to neural activity

**SUMMARY:** Metabolic coordination between neurons and astrocytes is critical for the health of the brain. However, neuron-astrocyte coupling of lipid metabolism, particularly in response to neural activity, remains largely uncharacterized. Here, we demonstrate that toxic, peroxidated fatty acids (FAs) produced in hyperactive neurons are transferred to astrocytic lipid droplets by lipoprotein particles. Astrocytes consume the FAs stored in lipid droplets via mitochondrial β-oxidation in response to neuronal activity and turn on a detoxification gene expression program. Together, our findings reveal that FA metabolism between neurons and astrocytes is coupled in an activity-dependent manner to protect neurons from FA toxicity. This coordinated mechanism for metabolizing FAs could underlie both homeostasis and a variety of disease states of the brain.

## INTRODUCTION

Neurons and astrocytes operate as a tightly coupled unit for energy metabolism in the brain. While neurons consume a considerable amount of ATP on the recovery of ion gradients challenged by postsynaptic potentials, action potentials and neurotransmitter recycling, astrocytes serve as the metabolic workhorses. Specifically, astrocytes provide neurons with metabolic substrates and antioxidants (Belanger et al., 2011; Belanger and Magistretti, 2009). This metabolic support allows neurons to allocate more cellular resources to sustain high activity rates during information processing.

An important and understudied area in neuron-astrocyte interaction relates to the metabolism of fatty acids (FAs). FAs are components of phospholipids in cellular membranes and are stored within cells as energy-rich triacylglycerides localized in lipid droplets (LDs). Storage of FAs in LDs serves two purposes. First, it removes from the cytoplasm excess free FAs, which are toxic and disrupt mitochondrial membrane integrity (Unger et al., 2010; Nguyen et al., 2017). Second, LD storage of FAs provides a specialized conduit for delivering FAs into mitochondria for their consumption as an alternative energy source during periods of nutrient depletion and stress (Rambold et al., 2015). Interestingly, neurons do not typically make LDs and have a low capacity for FA consumption in mitochondria for energy production (Schonfeld and Reiser, 2013). This makes neurons particularly sensitive to conditions of continued neuronal stimulation, in which high ROS levels induce peroxidation of FAs (Reynolds and Hastings, 1995). Unless they can destroy or remove these peroxidated FAs, neurons that are continually stimulated will undergo pathophysiology, giving rise to neurodegeneration (Sultana et al., 2013).

Unlike neurons, astrocytes make LDs, as well as other anti-oxidants, allowing them to effectively manage oxidative stress (Belanger and Magistretti, 2009). Given the tight coordination in energy metabolism between neurons and astrocytes, the question arises as to whether astrocytes help highly active neurons avoid FA toxicity from ROS buildup, for example, by taking up damaged FAs from neurons and storing them in LDs. Consistent with this possibility, recent studies have shown that oxidative stress in neurons triggers LD formation in neighboring astrocytes (Liu et al., 2015; Bailey et al., 2015), and that this is dependent on the presence of apolipoproteins (Liu et al., 2017). This suggests that stimulated neurons transfer their damaged FAs to astrocytes via lipoprotein particles, but how this occurs and the functional response(s) by astrocytes to this transfer has not been characterized.

Here, we investigate the cellular mechanisms of neuron-astrocyte metabolic coupling that protects neurons from FA toxicity during periods of neuronal stimulation. We first demonstrate that neuronal FAs are peroxidated during continued neuronal stimulation. Rather than storing these toxic FAs in LDs, we show that stimulated neurons expel them in association with lipoprotein-like particles secreted by neurons and/or astrocytes. Nearby astrocytes then take up these particles by endocytosis and store the peroxidated FAs in LDs. We also report that astrocytes respond to increased neuronal stimulation by enhancing breakdown of LDs and feeding liberated FAs into the mitochondria for use as fuel for oxidative phosphorylation. During this process, astrocytes upregulate genes involved in neutralizing oxidative species such as lipid peroxides, as well as genes involved in energy metabolism. Astrocytes thus help protect neurons during excessive stimulation both by taking up neuronal-released FAs and switching on genes for FA detoxification and FA-based oxidative energy metabolism. The activity-dependent stimulation of lipid metabolism by astrocytes provides a program for avoiding FA toxicity in response to the fluctuating needs of neurons.

## RESULTS

### Effect of excitotoxicity on the systems that regulate FA trafficking in neurons

To explore the trafficking itinerary of FAs in highly active neurons, we stimulated cultured hippocampal neurons with NMDA to induce excitotoxicity. Specifically, NMDA treatment is predicted to peroxidate lipids and FAs (Haba et al., 1991; Parihar and Hemnani, 2004), leading neurons to respond through potential changes in autophagy, LD biogenesis and mitochondria fusion dynamics. To assess whether lipids undergo peroxidation following NMDA treatment, we incubated neurons with linoleamide alkyne (LAA), which passively incorporates into cellular membranes. Upon lipid peroxidation, reactive aldehydes are produced which modify proteins that can be detected using Alexa Fluor 488-conjugated Click-iT chemistry (Figure 1A). We observed increased Alexa Fluor 488 fluorescence in neurons following NMDA treatment compared to control cells (Figure 1B-1C and S1A), supporting the view that excitotoxicity through NMDA treatment increases neuronal lipid peroxidation in hippocampal neuronal cultures.

**Figure 1.**
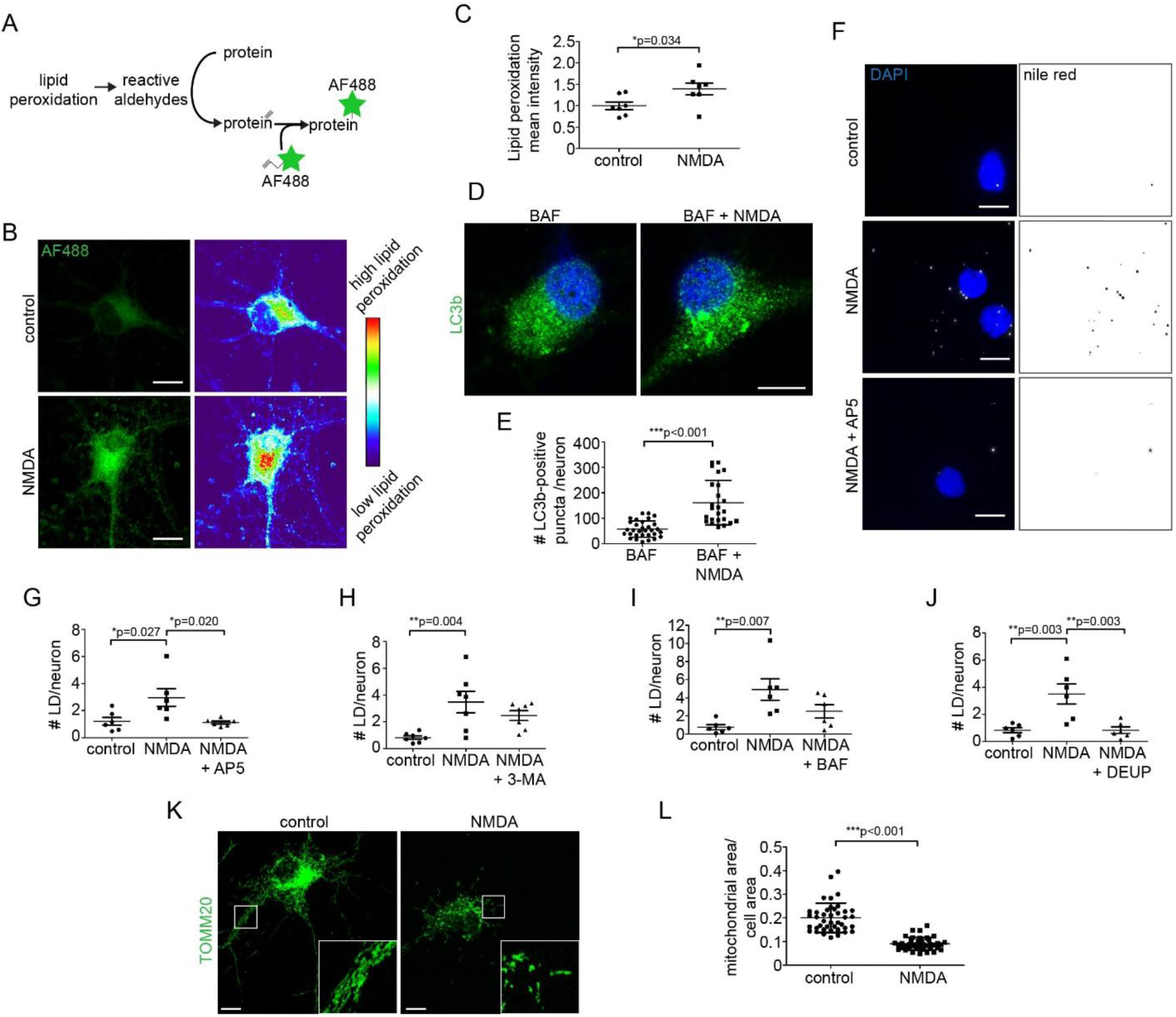
Excitotoxicity as an oxidative stress in neurons. (A) Schematic of Click-iT lipid peroxidation assay. (B-C) Neurons +/− NMDA were assayed for lipid peroxidation. Three independent experiments; n = 7 coverslips/treatment; 17.2 +/− 5.4 cells cells/coverslip; mean +/− SEM. (D-E) Neurons +/− NMDA were immunostained for LC3b. Two independent experiments; n = 3 coverslips/treatment; 15.7 +/− 7.0 cells/coverslip; mean +/− SEM. (F-G) Neurons +/− NMDA and AP5 were stained with Nile Red. Maximum intensity projections are displayed. Five independent experiments; n = 6 coverslips/treatment; 26.4 +/− 5.9 cells/coverslip; mean +/− SEM. (H) Neurons +/− NMDA and 3-MA were stained with BD493. Four independent experiments; n = 7 coverslips/treatment; 25.2 +/− 4.5 cells/coverslip; mean +/− SEM. (I) Neurons +/− NMDA and BAF were stained with BD493. Three independent experiments; n = 6 coverslips/treatment; 33.8 +/− 7.4 cells/coverslip; mean +/− SEM. (J) Neurons +/− NMDA and DEUP were stained with BD493. Three independent experiments; n = 6 coverslips/treatment; 30.5 +/− 4.2 cells/coverslip mean +/− SEM. (K-L) Neurons +/− NMDA were immunostained with TOMM20-AF488. Two independent experiments; n = 60 cells/treatment; mean +/− SD. Scale bars are 10 μm.

Neurons that have acquired peroxidated lipids in their membranes under excitotoxicity need a way to remove them. One way is through autophagy, the process whereby portions of an organelle or membrane are delivered to the lysosome for breakdown into simpler components. We therefore tested whether excitotoxicity through NMDA treatment caused an increase in autophagy in our hippocampal neurons using the autophagy marker LC3. This was accomplished using bafilomycin to prevent autophagosome fusion with lysosomes (Redmann et al., 2017), thereby allowing autophagosomes to accumulate within cells. When neurons were stimulated with NMDA in the presence of bafilomycin, increased levels of the LC3b were detected, indicating excitotoxicity enhances neuronal autophagy (Figures 1D and 1E). This result is consistent with peroxidated lipids in membranes of excited neurons being mobilized by autophagy for eventual removal from these cells.

Peroxidated lipids delivered to lysosomes through autophagy are expected to be broken down into FAs, which can quickly leak into the cytoplasm. To avoid FA toxicity from FA accumulation in the cytoplasm, autophagy-mobilized FAs in non-neuronal cells are efficiently delivered to LDs, increasing their number (Unger et al., 2010; Rambold et al., 2015). We tested, therefore, whether LDs were induced in NMDA-treated neurons. We first confirmed the neurons could form LDs by incubating with oleic acid, a long chain monounsaturated FA. A massive accumulation of LDs was seen (Figures S1B and S1C). Next, we tested if neurons form LDs in response to NMDA-induced excitotoxicity. A modest but significant increase in the number of LDs was observed (Figures 1F-G and S1D-E), which could be prevented with the NMDA receptor antagonist AP5 (Figures 1F and 1G).

Autophagy was responsible for this increase in LD formation following neuronal excitotoxicity since NMDA-induced LD formation was reduced by blocking autophagosome formation using 3-methyladenine (3-MA) (Figure 1H) or by inhibiting lysosomal autophagosome degradation using bafilomycin A1 (Figure 1I). We also observed a reduction in NMDA-induced LD formation in the presence of the pan-lipase inhibitor diethylumbelliferyl phosphate (DEUP) (Figure 1J), which taken together with involvement of autophagy points to the role of lysosomal lipases in the LD formation. Collectively, these data suggested that following excitotoxicity, neurons generate excess FAs by autophagy and some of these FAs are stored in a small number of LDs.

We wondered why there were not more LDs forming in cells undergoing NMDA-treatment. One possibility is that FAs are being efficiently delivered from LDs to mitochondria and broken down there through mitochondrial β-oxidation (Rambold et al., 2015). However, prior work has shown that neurons have a low capacity for FA degradation in mitochondria (Schonfeld and Reiser, 2017). Moreover, to efficiently degrade FAs in this manner the mitochondria network would have to be extensively fused (Rambold et al., 2015). We examined, therefore, mitochondrial morphology and quantified total mitochondrial area in control neurons and neurons treated with NMDA to evaluate mitochondrial capacity to degrade FAs. Rather than being extensively fused, the mitochondria present in NMDA-treated neurons were highly fragmented (Figure 1K and S1F) (Nguyen et al., 2011). Additionally, there was a decrease in mitochondrial area in neurons treated with NMDA (Figure 1L), suggesting mitochondria were being consumed by the increased autophagy occurring in these cells. Consistent with the remaining fragmented mitochondria being unable to consume FAs, we preloaded neurons with Red-C12 and were unable to detect the labelled FA in the mitochondria following NMDA treatment (Figure S1G). Based on these results, we concluded that excess, free FAs generated by autophagy in neurons treated with NMDA were not being turned over within mitochondria. Rather, some other pathway was removing these FAs from neurons to avoid FA toxicity.

### Highly active neurons release FAs in dense carriers

In non-neuronal cells containing highly fragmented mitochondria, FAs are expulsed and taken-up by neighboring cells with non-fragmented mitochondria (Rambold et al., 2015). We thus tested whether FAs in neurons could be released and taken up by neighboring astrocytes. To do so, we developed a transfer assay to monitor FA trafficking using BODIPY 558/568 C12 (Red-C12), a fluorescently labelled, saturated FA analogue (Rambold et al., 2015; Wang et al., 2010). We first incubated neurons with trace amounts of Red-C12 overnight, then co-cultured the labelled neurons with astrocytes on different coverslips separated by paraffin wax (Figure 2A). We found that fluorescently labelled FAs in neurons are transferred to GFAP-positive astrocytes in the absence of cell contact and that these FAs accumulate in LDs (Figure 2A-C, S2A-B). To exclude the possibility that contaminating astrocytes in our neuronal cultures were responsible for transferring the FAs, we performed transfer assays with neuronal cultures grown in Ara-C to eliminate proliferating glial cells. We found that neuronal cultures with over 98% purity transfer labelled FAs to astrocytes (Figure S2C-F). Therefore, neurons transfer FAs to neighboring astrocytes *in vitro*.

**Figure 2.**
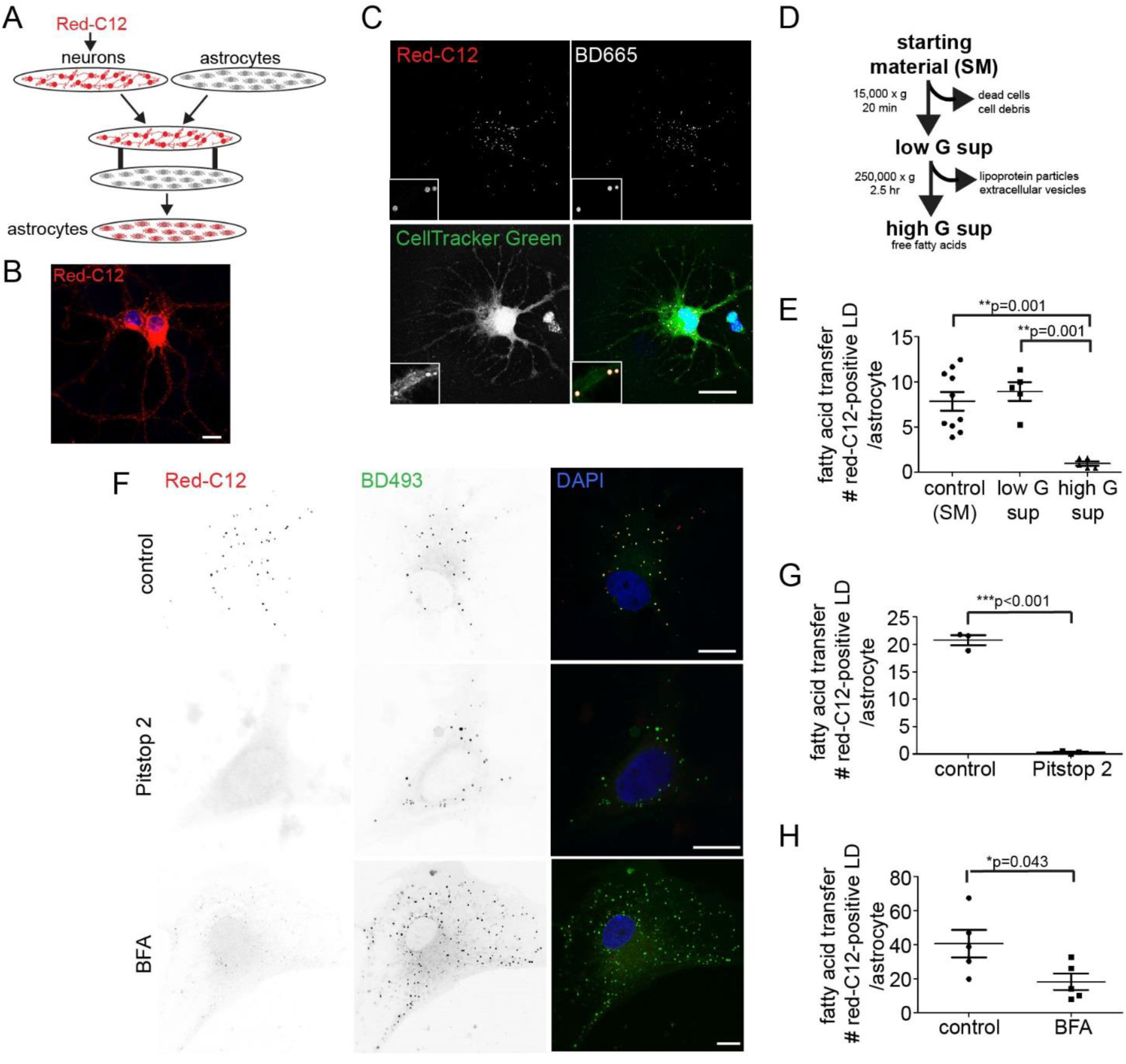
Neurons transfer FA to astrocytes. (A) Schematic of FA transfer assay. Neurons were incubated with Red-C12 overnight and incubated with astrocytes on separate coverslips. (B) Neurons were incubated with 2 μM Red-C12 overnight, chased for 1 hour in complete media, fixed and imaged. (C) Astrocytes analyzed for Red-C12 in LDs labelled with BD665. Astrocytes labelled using CellTracker Green. (D) Schematic of assay. Neuron-conditioned media was depleted of dead cells and debris by low-speed centrifugation and of lipoprotein particles by high-speed centrifugation. The resulting supernatant (sup) was used to treat astrocytes. (E) Neuron-conditioned aCSF was centrifuged and used to treat astrocytes. Astrocytes were fixed and analyzed for the appearance of Red-C12. Three independent experiments; minimum of n = 5 coverslips/treatment; 23.5 +/− 10.0 cells/coverslip; mean +/− SEM. (F) Red-C12 pulse chase assay described in A in the presence of DMSO, Pitstop 2 or Brefeldin A (BFA). Astrocytes were analyzed for the appearance of Red-C12. (G) Quantification of F. Two independent experiments; n = 3 coverslips/treatment. 23.5 +/− 3.5 cells/coverslip; mean +/− SEM. (H) Quantification of F. Four independent experiments; n = 6 coverslips/treatment. 27.5 +/− 3.9 cells; mean +/− SEM. All images are maximum intensity projections. Scale bars are 10 μm.

We next sought to determine the mechanism of this FA transfer. We took advantage of differential centrifugation to deplete the various components from the media of neurons labelled with Red-C12 (Figure 2D). We found that following low speed centrifugation to remove dead cells and debris, neuron-conditioned media continues to supply labelled FAs to astrocytic LDs (Figure 2E). Consistent with this, we observed that astrocytic LDs were labeled with Red-C12 following incubation with 0.2 μm pore filtered neuron-conditioned media (Figure S2G). These results suggested that FAs are being transferred as free FAs or as small dense carriers such as lipoprotein particles from neurons to astrocytes.

Following high-speed centrifugation to remove small dense carriers, the transfer of labelled FAs from neuron-conditioned media to astrocytes was eliminated (Figure 2E and S2H). As free FAs such as BSA-oleic acid remain in the supernatant following high speed centrifugation (Figure S2I), these data pointed to small dense carriers as the vehicle for FA transfer from neurons to astrocytes. Consistent with this model, FA transfer to astrocytes was blocked by inhibiting endocytosis with Pitstop 2, suggesting that internalization of small dense carriers by astrocytes is required for the transfer (Figure 2F and 2G). Furthermore, transfer of FAs was reduced by the addition of brefeldin A (BFA), a fungal metabolite that blocks export from the endoplasmic reticulum into the secretory pathway (Klausner et al., 1992)(Figure 2F and 2H). As BFA does not block endocytosis (Lippincott-Schwartz et al., 1991), this suggested that neuronal FA release and association with small dense carriers was dependent on the secretory pathway.

### Astrocytes internalize FAs secreted by neurons in lipoprotein particles

ApoE, a component of lipoprotein particles, is required for astrocytic LD accumulation in response to oxidative stress (Liu *et al.*, 2017). To test the hypothesis that lipoprotein particles might mediate transfer of FAs from neurons to astrocytes, we added excess purified LDL or oxidatively-modified oxLDL in a pulse-chase assay to competitively block lipoprotein particles receptors on astrocytes. We found that incubation with LDL or oxLDL dramatically reduced Red-C12 transfer from neurons to astrocytes (Figure 3A), indicating that internalization of lipoprotein particles is important for astrocytes to receive FAs. Furthermore, astrocytes increase LD number when treated with oxLDL containing lipid peroxides and their degradation products (Parthasarathy et al., 2010), but not LDL. (Figure 3B). On the other hand, neither LDL nor oxLDL induce LD formation in neurons (Figure S3A). This data suggested that astrocytes likely play an important role in internalizing and managing damaged lipid products from circulating lipoprotein particles in the brain.

**Figure 3.**
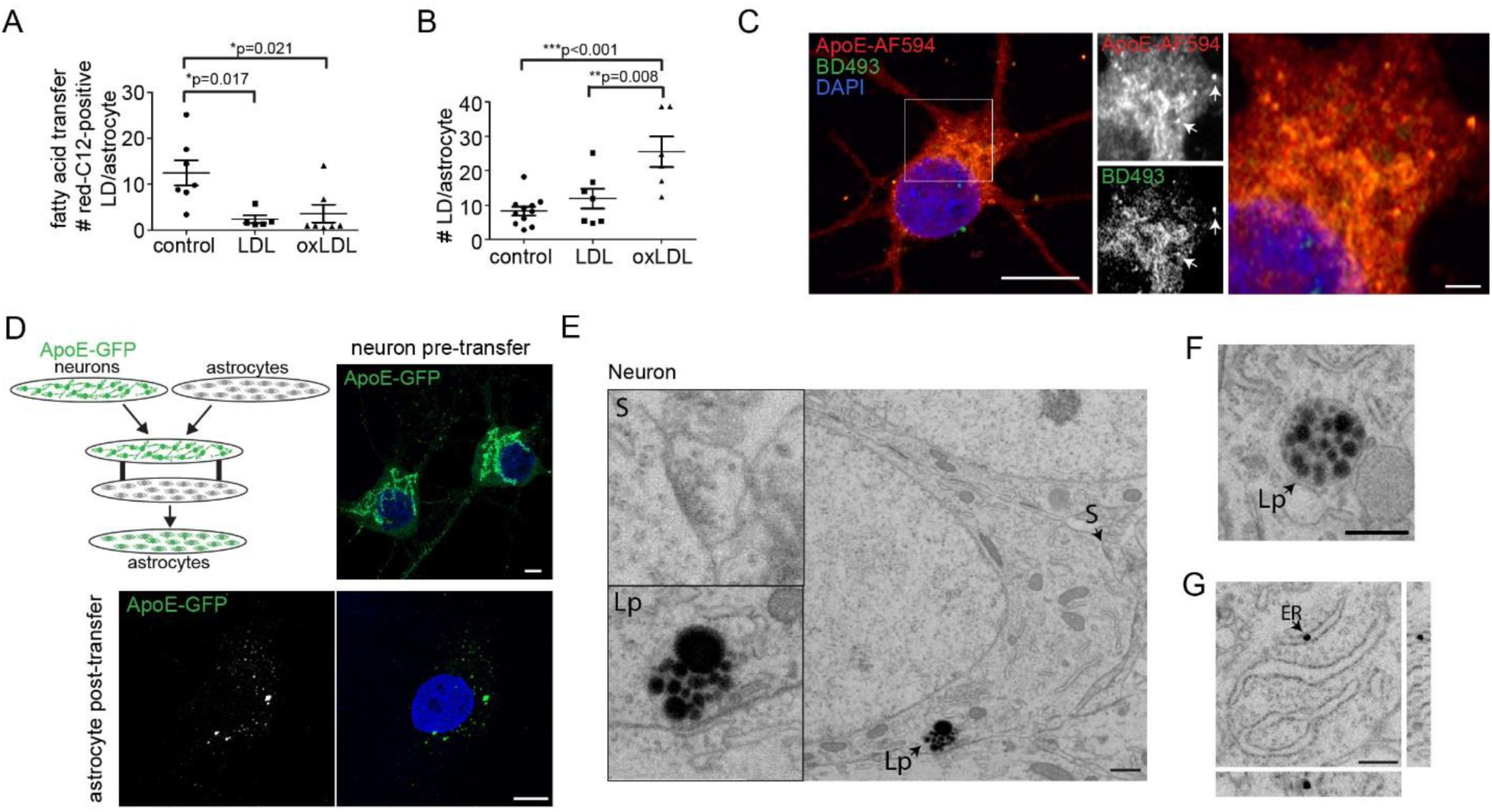
Neurons transfer FA by lipoprotein particles. (A) FA transfer assay as described in Figure 2A in the presence of LDL or oxLDL. Astrocytes were fixed and analyzed for the appearance of Red-C12. Three independent experiments; minimum of n = 5 coverslips/treatment; 18.2 +/− 3.4 cells; mean +/− SEM. (B) Astrocytes were treated with LDL or oxLDL and stained with BD493. Three independent experiments; minimum of n = 6 coverslips/treatment. 23.7 +/− 4.3 cells/coverslip; mean +/− SEM. (C) Neurons were immunostained with anti-ApoE and BD493. Boxed area magnified in right panels. Arrows highlight co-localization. Scale bars are 10 μm and 1 μm. (D) Top left panel: schematic of ApoE transfer assay. Top right panel: neurons expressing GFP-ApoE. Bottom Panel: Astrocytes post-transfer assay were analyzed for the appearance of ApoE-GFP. Scale bars are 10 μm. (E-G) Pyramidal neuron from CA1 in hippocampus was imaged by FIB-SEM. Lp indicates to lipid particles in membrane-enclosed vesicles, magnified on right. S indicates synapse. ER indicates lipid-dense structure in lumen of the ER. Orthogonal view of ER structure included. Scale bars are 500 nm.

We next asked what role neurons may play in regulating lipoprotein particles. Neurons express ApoE in cultured neurons (Figure 3C) (Dekroon and Armati, 2001) and following excitotoxicity *in vivo* (Xu et al., 2006). We reasoned that ApoE secreted by neurons (Figure S2H) could associate with lipoprotein particles and be internalized by astrocytes. To address this, we tested whether neuron-derived ApoE-GFP is transferred to astrocytes by performing a modified transfer assay. We found that neurons transferred ApoE-GFP to astrocytes (Figures 3D), confirming both that neurons secrete ApoE and that the secreted ApoE can be taken up by astrocytes. Neurons are thought to release non-lipidated ApoE that associates with astrocyte-derived lipoprotein particles (DeMattos et al., 1998). However, we observed endogenous ApoE in cultured neurons co-localizing with BD493, a neutral lipid stain, in small puncta and on Golgi-like structures (Figure 3C). This suggested that neurons are releasing lipidated ApoE and forming lipoprotein-like particles.

To better assess whether neurons can make lipoprotein-like particles, we employed focused ion beam scanning electron microscopy (FIB-SEM) technology (Xu et al., 2017), a transformative technology that allows 3D isotropic imaging of cells and tissues at nanometric resolution. We examined a volume of 60 × 80 × 45 μm^3^ with 8 × 8 × 8 nm^3^ voxel resolution from an adult mouse hippocampus. The hippocampal tissue was prepared with imidazole-buffered osmium tetroxide to stain lipids such as LDs and lipoprotein particles (Zhang et al., 2011; Rong et al., 2015). In single 8 nm thick planes, we observed membrane-bound lipid-dense structures in neurons (identified by the presence of synapses) in the pyramidal cell layer (Figure 3E-F and Movie S1). These structures were distinguishable from autophagosomes by their absence of organellar components (Figure S3B) and resembled lipoprotein particles (Rong et al., 2015). Similar lipid-dense, particle-like structures were also seen in the ER (Figure 3F and S3D), where lipid-particles are known to be synthesized. Collectively, these results support the idea that neurons can synthesize lipid-rich particles in the ER and secrete them, opening the way for astrocytes to take them up by endocytosis. Assuming neuronal-derived FAs can attach to these lipid-rich particles, either in the ER or after the particles are secreted (through FA transporters), the result would be an extracellular pool of FA-laden lipid particles available for uptake by astrocytes.

### LD formation in astrocytes following acute stroke

Our results showing the transfer of FAs from neurons to astrocytes *in vitro* led us to ask whether this occurs *in vivo*, especially under conditions of oxidative stress. To address this, we performed a pial strip lesion in the rat brain cortex, which has been used as a model system of acute stroke (Farr and Whishaw, 2002; Alaverdashvili et al., 2008). The assay entails de-vascularizing a region of the motor cortex by removal of the pia mater. We then examined whether there was a buildup of LDs in astrocytes, which we used as a readout of FA transfer from neurons. We first tested the efficacy of the assay by immunolabeling with markers of oxidative injury. A major source of tissue damage following stroke is excitotoxity caused by excess glutamate release (Lai *et al.*, 2014). Consistent with this, we observed enhanced expression of the immediate early gene c-fos, a marker of neuronal activation, in the stroke region compared to the contralateral hemisphere (Figure 4A and 4B). Following the brain injury, astrocytes also upregulate expression of the intermediate filament protein GFAP and microglia migrate to the region of damage (Ren et al., 2013; Ben et al., 2015). We found an accumulation of GFAP-positive astrocytes and Iba1-positive microglia in the lesion site compared to the contralateral hemisphere (Figure S4A). Together these data confirm that our acute stroke model was sufficient to evoke oxidative injury.

**Figure 4.**
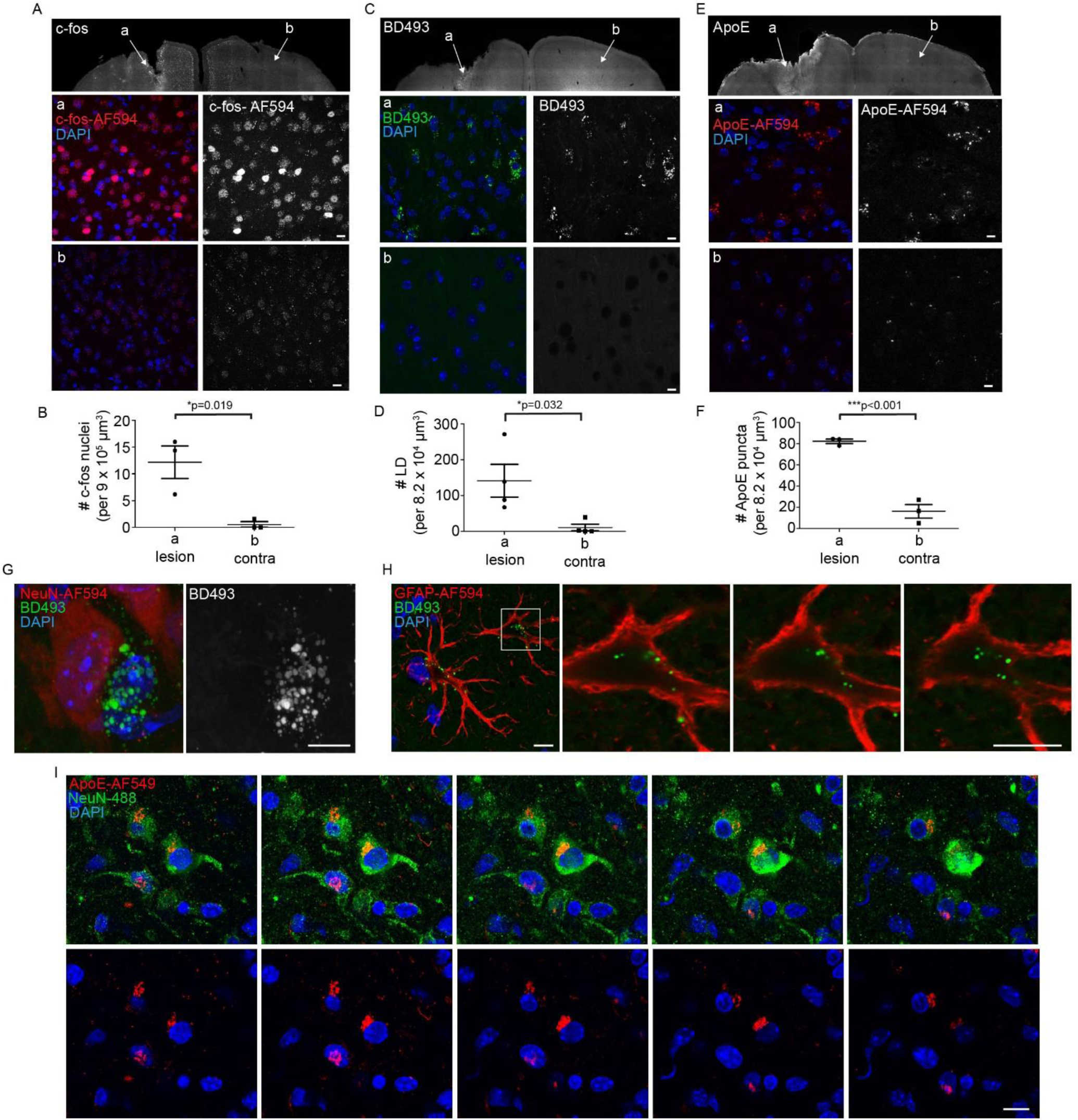
Astrocytes accumulate LDs following acute stroke. (A-F) Mouse cortex post pial strip lesion was stained with BD493, c-fos or ApoE. Bottom panels are magnified maximum intensity projections in the (a) lesioned hemisphere and (b) contralateral hemisphere. Scale bars are 10 μm. n = 4 animals for BD493 and n=3 animals for c-fos and ApoE; mean +/− SEM. (G-H) Mouse cortex post pial strip lesion was immunostained with anti-NeuN or GFAP. Maximum intensity projections. Boxed area is magnified in right panels. Scale bars are 10 μm. (I) Mouse cortex post pial strip lesion was immunostained with anti-NeuN and ApoE and imaged in 0.5 μm sections. Scale bars are 10 μm.

We next examined whether acute stroke induced LD formation. As predicted, we observed LD accumulation in the lesion site but not in the contralateral hemisphere following pial strip lesion (Figure 4C and 4D). We were unable to detect LDs in neurons following stroke (Figure 4G), consistent with our *in vitro* assays showing that neurons transfer FAs to astrocytes (Figure 2 and 3). Instead, we found LD accumulation within the soma and along the processes of neighboring GFAP-positive astrocytes and Iba1-positive microglia (Figure 4H and S4B).

To test whether lipoprotein particle-mediated FA transfer was involved in the accumulation of astrocytic LDs, we immunolabelled the tissue for ApoE. An increase in ApoE staining was observed in the lesion site as compared to the contralateral hemisphere (Figure 4E and 4F). In fact, a portion of neurons were found to express ApoE following acute stroke *in vivo* (Figure 4I). Together, these data support the role of lipoprotein particle-mediated transfer of FAs from neurons to astrocytes following oxidative injury in the form of acute stroke.

### Neural activity stimulates FA transport from neurons to astrocytes

Since stroke involves oxidative damage to both neurons and astrocytes (Lai *et al.*, 2014), we next sought to test whether hyperactivity in the neurons alone was sufficient to trigger FA transfer and LD formation in astrocytes *in vivo*. To address this question, we used adeno-associated virus to express the chemogenetic receptor hM3Dq-mCh under the neuron specific promoters (synapsin or CamKII) in the motor cortex of mice (Figure 5A and 5B). Intraperitoneal injection of the inert ligand clozapine-N-oxide (CNO) activates neurons expressing the hM3Dq receptor (Alexander et al., 2009). We stimulated neurons over 3 days and confirmed elevated neuronal activity by observing an upregulation of c-fos in neurons expressing hM3Dq treated with CNO, but not in those treated with saline, or CNO-treated neurons not expressing hM3Dq in the contralateral hemisphere (Figure 5C and 5D). We also observed an accumulation of LDs in the injected region after treatment with CNO (Figure 5E and 5F). Similar to the stroke model, we found LD accumulation in astrocytes but not in neurons (Figure 5G and 5H). These results strongly suggest that neurons transfer FAs to astrocytic LDs in response to enhanced neural activity *in vivo*.

**Figure 5.**
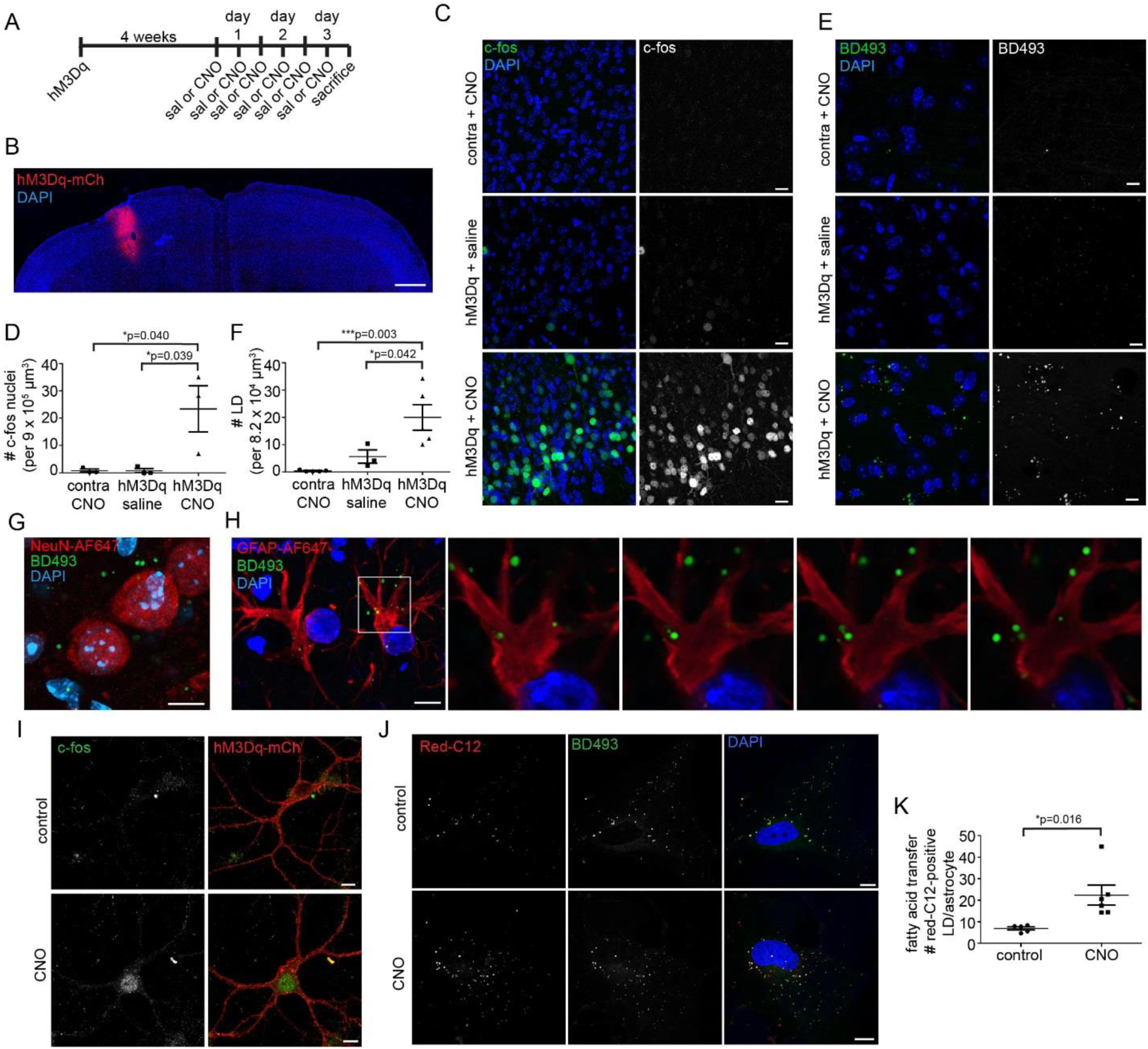
Neural activity stimulates FA transfer to astrocytes. (A) Schematic of *in vivo* activation assay. HM3Dq-mCh under neuron specific promotors Syn or CamKII was injected in the motor cortex. Neurons were stimulated with intraperitoneal injection of clozapine-N-oxide (CNO) for 3 days. Control animals were injected with saline (Sal). (B) Motor cortex expressing hM3Dq-mCh. Scale bars, 500 μm. (C-D) Motor cortex following neuronal activation was immunostained with anti-c-fos. Maximum intensity projection. Scale bars, 20 μm. n = 3 animals/treatment; mean +/− SEM. (E-F) Motor cortex following neuronal activation was stained with BD493. Maximum intensity projection. Scale bars, 10 μm. n = 3 animals/treatment; mean +/− SEM. (G-H) Motor cortex in hM3Dq-mCh expressing region following CNO treatment were immunostained with anti-NeuN and anti-GFAP. Maximum intensity projections. Boxed area is magnified in right panels. Scale bars,10 μm. (I) Cultured neuron expressing hM3Gq-mCh was treated with CNO for 90 minutes, fixed and immunostained with anti-c-fos. Scale bars, 10 μm. (J-K) FA transfer assay as described in Figure 2A using neurons expressing hM3Dq-mCh with or without CNO. Astrocytes were analyzed for the appearance of Red-C12-positive in BD493-positive LDs. Three independent experiments; minimum of n = 5 coverslips/treatment; 27.1 +/− 4.4 cells/coverslip; mean +/− SEM.

We next tested whether neuronal activity enhanced FA transfer using our *in vitro* pulse-chase transfer assay. First, we used adeno-associated virus to express hM3Dq-mCh in cultured neurons and showed they could be activated in response to CNO as evidenced by increased c-fos staining (Figure 5I). Neurons were preloaded with the labelled FA, Red-C12, and co-cultured with astrocytes on different coverslips in the presence or absence of CNO. We observed enhanced FA transfer upon neuronal stimulation by CNO (Figure 5J and 5K). Red-C12 transfer, as opposed to hMD3q-mCh transfer, was confirmed by co-localization with BD493-positive LDs (Figure 5J). The increased transfer occurred without changes in neuronal apoptosis as assessed by DAPI nuclear staining (Figure S5A-B). Together, these results reveal that the transfer of FAs from neurons to astrocytes was significantly increased during periods of enhanced activity.

### Neuronal activity decreases LD number in astrocytes through increased lipolysis

Since neurons release FAs during periods of enhanced activity, and astrocytes sense neuronal activity through glutamate receptors such as the NMDA receptor (Anderson and Swanson, 2000; Dzamba et al., 2013), we wondered whether neuronal activity influences LD number in astrocytes. To assess this, we tested whether the glutamate treatment alters the number of LDs in cultured astrocytes. A reduction in the number of LDs in cultured astrocytes treated with glutamate was observed (Figure S6A). This response involved the NMDA receptor since a similar reduction in the number of LDs occurred in astrocytes treated with NMDA, and the effect could be eliminated with the addition of the NMDA receptor antagonist AP5 (Figure 6A-B).

**Figure 6.**
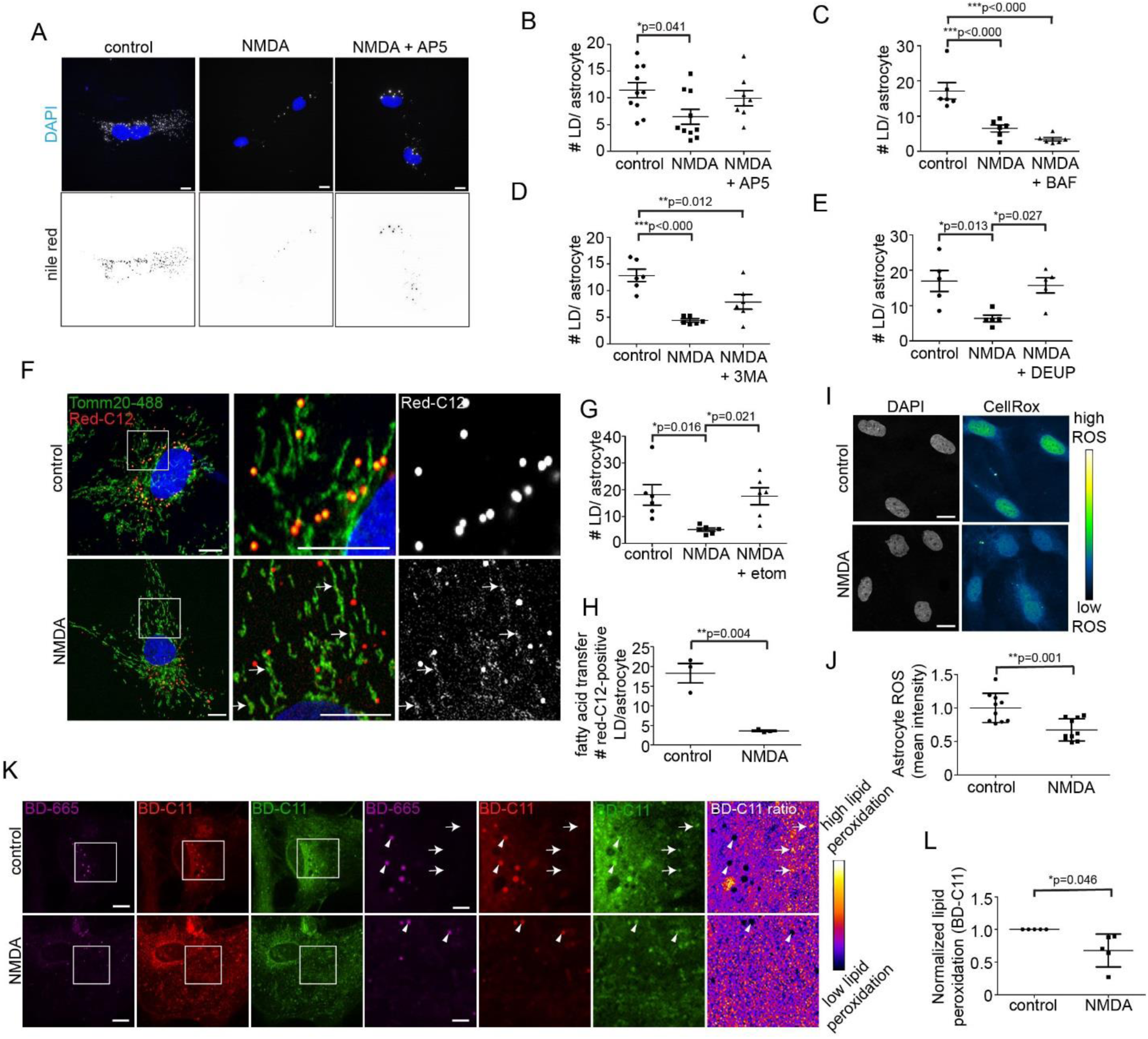
Astrocytes detoxify ROS and degrade FA in response to neuronal activity. (A-B) Astrocytes +/− NMDA and AP5 were stained with Nile Red. Five independent experiments; n = 8 coverslips/treatment; 21.2 +/− 4.5 cells/coverslip; mean +/− SEM. (C) Astrocytes +/− NMDA and BAF were stained with BD493. Three independent experiments; n = 6 coverslips/treatment; 26.0 +/− 5.3 cells/coverslip; mean +/− SEM. (D) Astrocytes +/− NMDA and 3MA were stained with BD493. Three independent experiments; n = 6 coverslips/treatment; 28.1 +/− 7.2 cells/coverslip; mean +/− SEM. (E) Astrocytes +/− NMDA and DEUP were stained with BD493. Three independent experiments; n = 5 coverslips/treatment; 24.3 +/− 6.5 cells/coverslip; mean +/− SEM. (F) Astrocytes were loaded with Red-C12 overnight, chased in +/− NMDA, fixed and immunostained with anti-TOMM20. Boxed area magnified on right. Scale bars,10 μm. (G) Astrocytes +/− NMDA and Etomoxir were stained with BD493. Three independent experiments; n = 6 coverslips/treatment; 28.5 +/− 4.1 cells/coverslip; mean +/− SEM. (H) FA transfer assay as described in Figure 2A +/− NMDA. Three independent experiments; n = 7 coverslips/treatment; 27.7 +/− 4.8 cells/coverslip; mean +/− SEM. (I-J) Astrocytes +/− NMDA were labelled with CellRox oxidative stress sensor. Four independent experiments; n = 7 coverslips/treatment; 29.5 +/− 5.2 cells/coverslip; mean +/− SEM. (K) Astrocytes +/− NMDA were labelled with BD665 and BD-C11. BD-C11 561 excitation (red) shows total lipid level while BD-C11 488 excitation (green) and ratio image shows peroxidated lipid. Boxed area highlighted on right. Arrows show peroxidated puncta. Arrowheads show low lipid peroxidation in LDs. Scale bars,10 μm. (L) Quantification of K. Five independent experiments; 18.9 +/− 5.7 cells/treatment normalized to control treatment to generate a mean +/− SD.

We speculated that the reduction in LD number in activated astrocytes was due to an increase in the rate of LD degradation, since this would allow the astrocyte to utilize the FAs in oxidative mitochondrial energy production. Astrocytes could utilize two possible mechanisms to break down LDs under NMDA receptor activation: lipophagy (the autophagic degradation of LDs), or lipolysis by cytoplasmic lipases. We found that autophagy inhibitors 3-MA and bafilomycin A1 had no effect on NMDA-dependent LD degradation (Figure 6C and 6D). By contrast, the pan-lipase inhibitor DEUP prevented NMDA-dependent LD degradation (Figure 6E). This indicated that cytoplasmic lipases rather than autophagic mechanisms were responsible for breaking down astrocytic LDs under increased neuronal activity. The data further confirmed our speculation that reduced LD numbers in activated astrocytes results from an increase in LD turnover in these cells.

### Astrocytes degrade FAs and neutralize ROS in response to neural activity

We next asked whether FAs liberated from LDs are degraded and consumed in astrocytes by FA oxidation in mitochondria. For this purpose, we preloaded astrocytes with Red-C12 and tracked the labelled FA after NMDA treatment to see if it was delivered to mitochondria. Consistent with this, Red-C12 appeared in mitochondria as early as 4 hours after NMDA treatment (Figure 6F). Co-treatment of astrocytes with NMDA and etomoxir, a carnitine palmitoyltransferase-1 inhibitor that prevents FA transport into mitochondria, decreased the NMDA-dependent reduction in the number of LDs (Figure 6G), suggesting astrocytes deliver FAs to mitochondria for β-oxidation in response to neuronal activity. We also observed a reduction in Red-C12 (originating from neurons) in astrocytes in our transfer assay when NMDA was added (Figure 6H), which would occur if astrocytes degrade neuron-derived FAs in response to neuronal activity. These results support a model in which neurons augment FA transport to astrocytes in response to neuronal activity (Figure 5), while astrocytes degrade the incoming FAs in response to that same activity.

Because mitochondrial β-oxidation itself can produce ROS (Schonfeld and Reiser, 2017), we reasoned that astrocytes with increased mitochondrial β-oxidation under conditions of excess neuronal activity must have a mechanism to neutralize ROS, otherwise they would jeopardize their own health. To examine this, we measured oxidative stress levels by staining astrocytes with the CellRox probe that fluoresces green upon oxidation by ROS in the nucleus. A decrease in CellRox fluorescence was observed in cultured astrocytes treated with NMDA, indicating decreased oxidative stress levels in these cells (Figure 6I, 6J and S6B). As a second test, we measured lipid peroxidation in astrocytes using the BODIPY-C11 ratiometric lipid peroxidation sensor. There was a decrease in levels of peroxidated lipids in both LDs and the surrounding membranes upon treatment of astrocytes with NMDA, indicating lipid peroxidation in astrocytes was reduced (Figure 6K and 6L). Taken together, these results suggest that under excess neuronal activity, astrocytes degrade incoming neuronal-derived FAs in mitochondria through β-oxidation, and this is associated with decreased ROS and lipid peroxidation levels in astrocytes.

### Astrocytes are equipped with the machinery to neutralize ROS

Since astrocytes internalize peroxidated lipids from neurons, we reasoned that astrocytes must be equipped with the molecular machinery to neutralize the oxidative stress and metabolize the lipids. One way this might occur is if astrocytes upregulate genes responsible for oxidative stress management in response to LD accumulation. To test this, we performed a gene expression profile of cultured astrocytes, in which the cells were analyzed based on whether or not they contained LDs. Astrocytes were stained with Nile Red to selectively label LDs under 488 nm excitation (Greenspan *et al.*, 1985). Cells containing LDs were then separated from those that did not by fluorescence activated cell sorting. RNA sequencing (RNAseq) analysis was then performed on both of the isolated cell populations (Figure S7A). By comparing the gene expression profile of our cultured astrocytes to an existing RNAseq database of cell types in the brain (Zhang *et al.*, 2014), we confirmed the purity of our astrocyte cultures, ruling out the possibility that contaminating cell types represented the cells that had LD buildup (Figure S7B). Notably, a group of genes related to neutralizing oxidative stress and lipid metabolism were upregulated in astrocytes containing LDs (Figure 7A-B and S7C).

**Figure 7.**
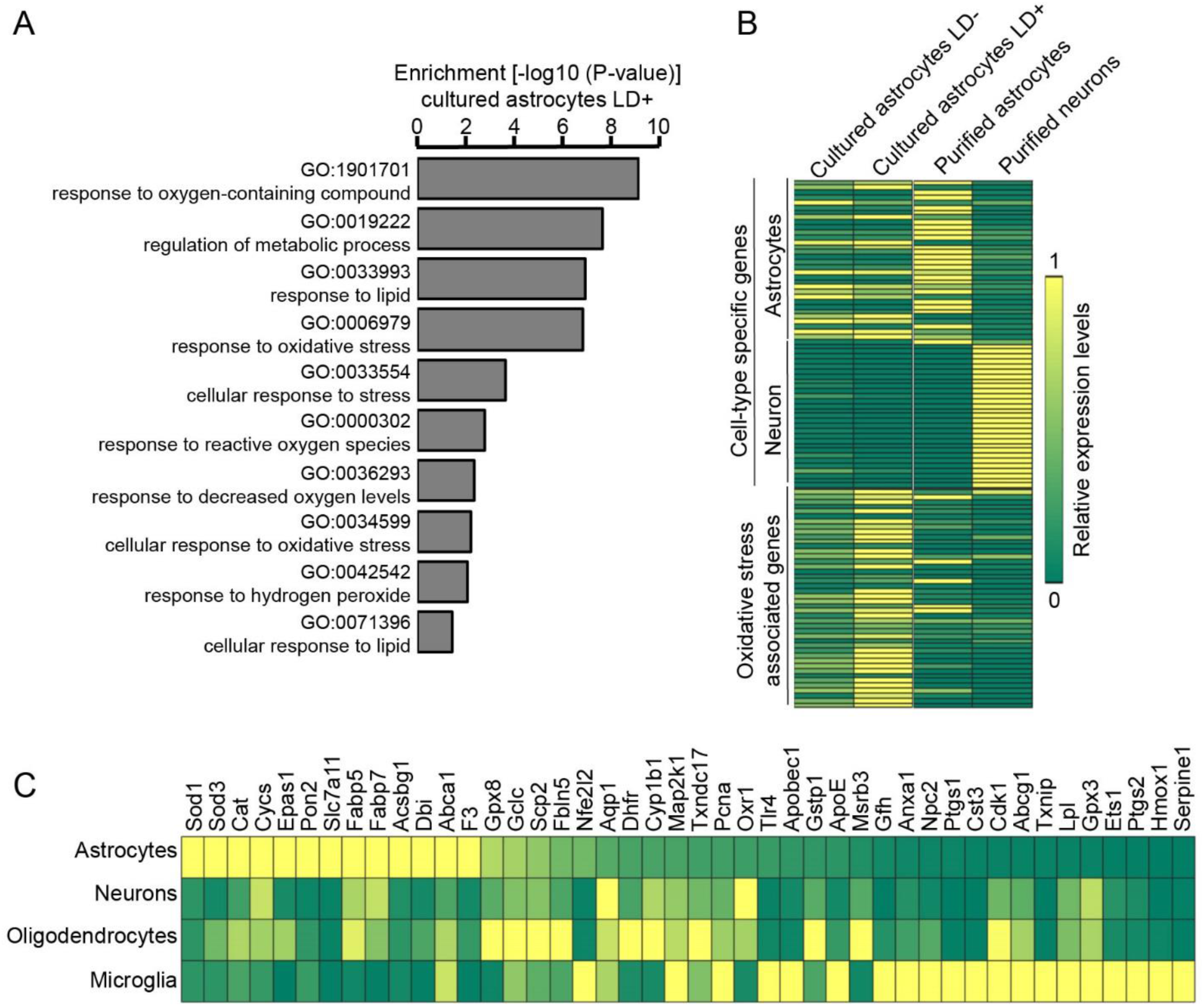
Up-regulation of genes associated with FA metabolism and oxidant detoxification in LD-positive astrocytes. (A) Gene Ontology (GO) pathways enriched in cultured astrocytes containing LDs. (B) Astrocytes enriched for genes associated with oxidative stress compared to neurons. Relative gene expression levels in different cell types were presented in the heat-map. (C) Expression levels of selected genes in purified cells. Genes selected as those associated with oxidative stress and enriched in astrocytes containing LDs in B. The color-bar is the same as in B.

Our differential gene expression analysis also revealed that astrocytes expressed higher levels of oxidative stress-and lipid metabolism-related genes compared to neurons, regardless of whether they were purified from the brain or cultured, or if they contained LDs (Figure 7B). Among the genes enriched in astrocytes containing LDs were those responsible for neutralizing oxidative species such as lipid peroxides (Gpx8), superoxide radicals (Sod1 and Sod3) and hydrogen peroxide (Cat) and genes involved in FA transport (Fabp5 and Fabp7) and FA metabolism (Acsbg1 and Dbi). Interestingly, RNAseq analysis of cells purified from the brain shows that while microglia and oligodendrocytes also express genes involved in responding to oxidative stress, they used distinct sets of genes from astrocytes and each other. Therefore, each glial cell type would appear to use a unique machinery to manage oxidative stress at basal levels. However, astrocytes upregulate these genes in response to LD accumulation (Figure 7B). Importantly, neurons expressed comparatively low levels of these genes, indicating they are not adequately equipped to handle oxidative stress and lipid accumulation (Figure 7B and C). This supports the reliance of neurons on neighboring glia, such as astrocytes, to metabolize excess peroxidated FAs and neutralize oxidative stress.

## DISCUSSION

The regulation of lipid metabolism is critical for avoiding FA toxicity during neuronal stimulation and ultimately for maintaining the health of the brain. Here, we discovered that avoidance of FA toxicity in neurons is achieved by the activity-dependent coordination between neurons and astrocytes in the metabolism of FAs. Specifically, hyperactive neurons release toxic FAs as lipoprotein particles that neighboring astrocytes take up. These transferred FAs are used as metabolic intermediates for increased mitochondrial oxidation and detoxification in the astrocyte in response to neural activity. As discussed below, we propose that this mechanism allows astrocytes to protect hyperactive neurons from FA toxicity. The increased ATP produced by FA oxidation in astrocytes could also trigger a negative feedback loop that, in turn, would suppress hyperactive neurons by triggering inhibitory interneuron pathways.

When neurons are hyperactive, several changes in their metabolism take place. One recently observed effect is an increase in the rate of glycolysis compared to oxidative metabolism (Yellen, 2018). Glycolysis involves fewer steps than mitochondrial oxidative metabolism and therefore may respond more quickly to produce ATP. Glycolysis also results in greater FA synthesis than that under mitochondrial oxidative metabolism. In fact, increased FA synthesis has previously been demonstrated in neurons with defective mitochondria (Liu et al., 2015). A crucial predicted outcome of enhanced glycolysis in the active neuron, therefore, is boosted FA synthesis. This, we believe, is the ‘Achilles’ heel’ for the hyperactive neuron: its accumulation of toxic FAs that cannot be stored effectively in LDs due to the neuron’s limited capacity to make LDs.

The accumulation of free FAs in hyperactive neurons are toxic not only because of their susceptibility to peroxidation. Recent work has shown that in LD-defective cells, FAs can be converted into acylcarnitines, which cause mitochondrial fragmentation/dysfunction and ROS production (Nguyen et al., 2017). This process is likely to have profound consequences in hyperactive neurons since dysfunctional mitochondria are unable to metabolize FAs and their ROS production causes peroxidation of lipids on membranes. This is further accentuated by FA release from peroxidated membranes undergoing autophagy, generating a runaway system of toxic FA buildup. To relieve this stress, the neuron would need a mechanism to expel its toxic FAs. Our findings support this model by showing that in hyperactive neurons: LD number only minimally increases; mitochondria undergo fragmentation; levels of lipid peroxidation and autophagy significantly increase; and, FAs are released from the cells.

We envision two potential, non-exclusive pathways for release of FAs from hyperactive neurons. The first involves direct transport of free FAs through ATP-binding cassette (ABC) transporters on the neuronal cell surface. Prior work has shown that such transfer can lead to loading of circulating lipoprotein particles outside the cell (Kim et al., 2007). A second possible pathway for FA release is by loading FAs into lipoprotein-like particles within the lumen of the ER in the neuron. The ER-localized lipoprotein-like particles are then packaged into membrane-bound secretory vesicles that target to the plasma membrane for release. Supporting this exit possibility, we observed lipid particles in the ER and secretory vesicles by electron microscopy in brain tissue sections with properties consistent with lipoprotein-like particles. Further work is needed to clarify the relative extent to which hyperactive neurons use the lipid particle release pathway, the ABC transporter-based pathway, or both in expelling excess FAs.

Once FAs are released extracellularly, only those associated with lipoprotein particles are subsequently taken up by astrocytes and used metabolically. This was demonstrated in our neuron-astrocyte FA transfer assays, in which labeled FAs in neurons were transferred into astrocytes exclusively through the intermediary of lipoprotein particles, including ApoE-positive particles. ApoE, a lipid transport protein associated with lipoprotein particles, has previously been shown to be necessary for the appearance of glial LDs (Liu et al., 2017). While predominantly synthesized by astrocytes, we show that ApoE expression is also upregulated in neurons in response to oxidative stress in the form of acute stroke *in vivo*. Notably, we also found that ApoE transfers from neurons to astrocytes *in vitro*. Together, these results suggest that neurons may upregulate ApoE expression under oxidative stress specifically to increase their FA efflux through the lipoprotein-based pathway.

Further evidence of a role of ApoE in FA efflux from neurons is the toxic effect of an ApoE4 polymorphism with reduced lipid binding and secretion capacity when it is expressed in neurons (Buttini et al., 2010; Mahley, 2016). This could be explained if neurons expressing ApoE4 do not release FAs from the cell as efficiently as wildtype ApoE, and as a result undergo greater lipotoxicity and/or damage by lipid peroxidation. Introducing the ApoE4 into astrocytes has no similar effect on brain toxicity (Buttini et al., 2010; Mahley, 2016). Significantly, ApoE4 is the leading risk factor for late onset Alzheimer’s disease and determines neuronal susceptibility to excitotoxicity (Aoki et al., 2003; Buttini et al., 2010). Apolipoproteins such as ApoE, therefore, may play a key role in supporting neuronal health by channeling FAs away from neurons to astrocytes.

The existence of two possible pathways for astrocytes to internalize FAs from lipoprotein particles raised the question whether there is an advantage of endocytosing the particles over releasing FAs extracellularly and internalizing them via FA transporters. One possible advantage would be efficiency; endocytosed lipoprotein particles would deliver a large bolus of FAs into the cell. Another significant advantage of this pathway is that it could minimize FA toxicity within the astrocyte from incoming peroxidated lipids. This is because endocytosed lipoprotein particles carrying peroxidated lipids would be enclosed in a membrane-bound endosome and FAs can be quickly sequestered within LDs after releasing from lipoproteins in lysosomes (Davies et al., 2000; Guo et al., 2009; Zeng et al., 2003). For example, it has previously been reported that peroxidated FAs within astrocytes become sequestered in LDs to protect the cell from further damage (Liu et al., 2015). Helping to explain this, our FA transfer experiments in neuron-astrocyte co-cultures revealed that labeled FAs in neurons are delivered into LDs of astrocytes. Additionally, our in vivo experiments examining the response to a pial-strip brain lesion showed astrocytes at the wound site are enriched LDs. These findings suggest astrocytes protect neurons by serving as a sink for excess peroxidated and non-peroxidated FAs packaged into LDs.

FAs stored in LDs do not necessarily have an end-fate there, but can also be used as an energy source for oxidative phosphorylation by β-oxidation in mitochondria. We found that when astrocytes are stimulated with NMDA, they break down their LDs via mitochondrial oxidative metabolism and have reduced ROS production. We demonstrated this in several ways, including showing that FAs move from LDs to mitochondria in NMDA-treated astrocytes, and that LD number and ROS levels in these cells decrease through a mechanism dependent on LD-to-mitochondria transfer. Why might this effect of NMDA on astrocytes be significant for understanding the neuronal response to stimulation? Astrocytes express several glutamate receptors including the NMDA receptor, which allow them to sense and respond to neuronal activity (Anderson and Swanson, 2000; Dzamba et al., 2013). Glutamate released by stimulated neurons would activate NMDA receptors on nearby astrocytes and trigger the consumption of FAs by β-oxidation in the astrocyte, as we observed, resulting in increased ATP production.

Interestingly, it has previously been shown that glutamate accompanying neural activity also triggers ATP release by astrocytes (Zhang et al., 2003). The ATP released by astrocytes activates interneurons, resulting in increased synaptic inhibition within intact hippocampal circuits (Bowser and Khakh, 2004; Kawamura et al., 2004). Thus, it is possible that hyperactive neurons transfer their FAs to astrocytes, which use them to boost ATP levels and increase the activity of local inhibitory interneurons. This increased interneuron activity would consequently provide elevated feed-back inhibitory transmission onto the hyperactive neurons, potentially relieving them of excitotoxicity. Such a feed-back loop from astrocytes to neurons could be an important mechanism for attenuating the activity of hyperactive neurons and protect them from the damage associated with excess FA accumulation. The reduced ROS and boosted oxidative phosphorylation observed in astrocytes in response to NMDA is likely an important part of this system. Indeed, we found that LD-containing astrocytes expressed higher levels of genes involved in metabolism and ROS detoxification, reflecting enhanced mitochondrial function. Earlier studies have shown that astrocyte activation is involved in neuronal potentiation and memory function (Aguado et al., 2002; Adamsky et al., 2018) so an area for further study is how synaptic plasticity may be linked to the coordination of neurons and astrocytes in the metabolism of FAs.

## Materials and Methods

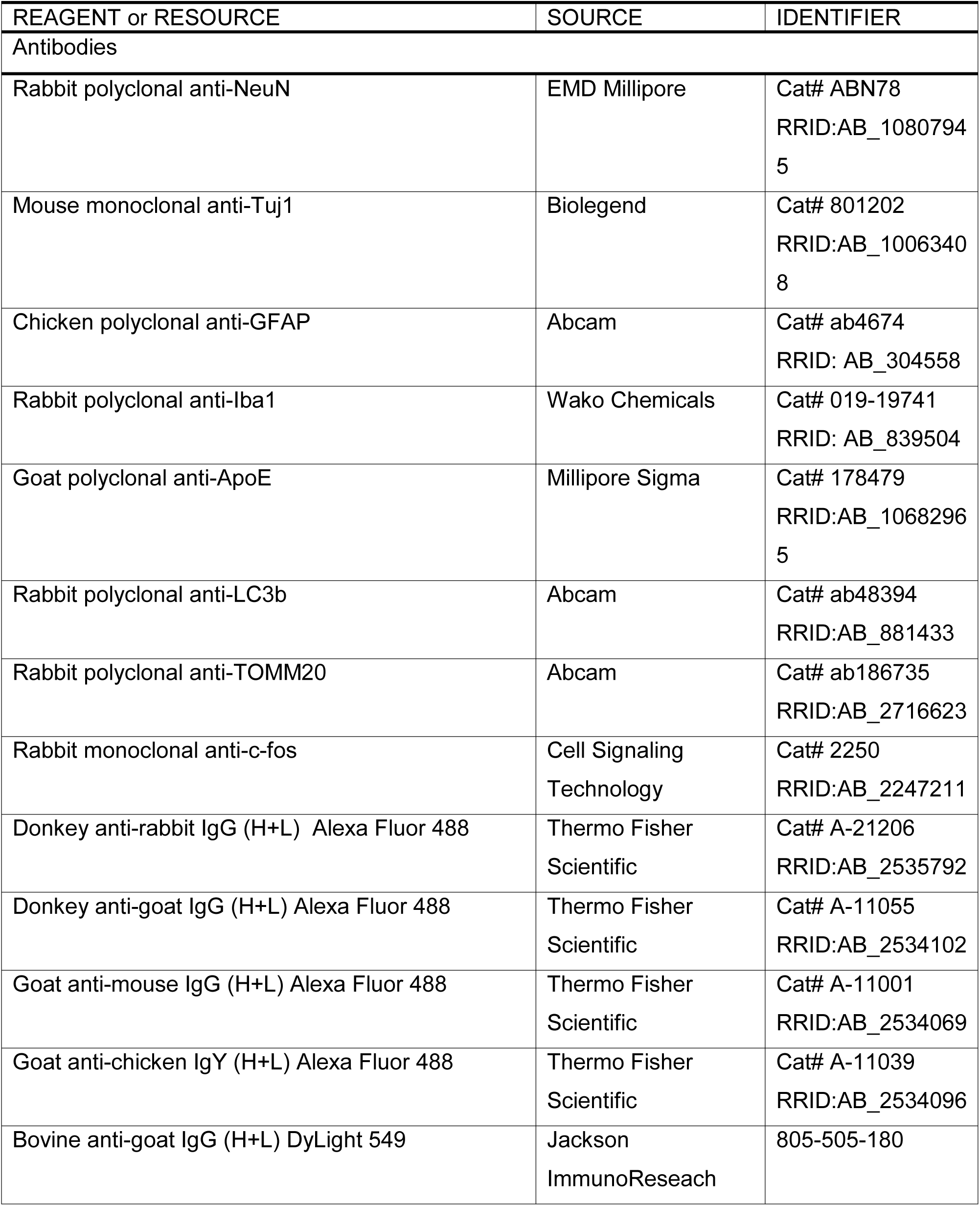
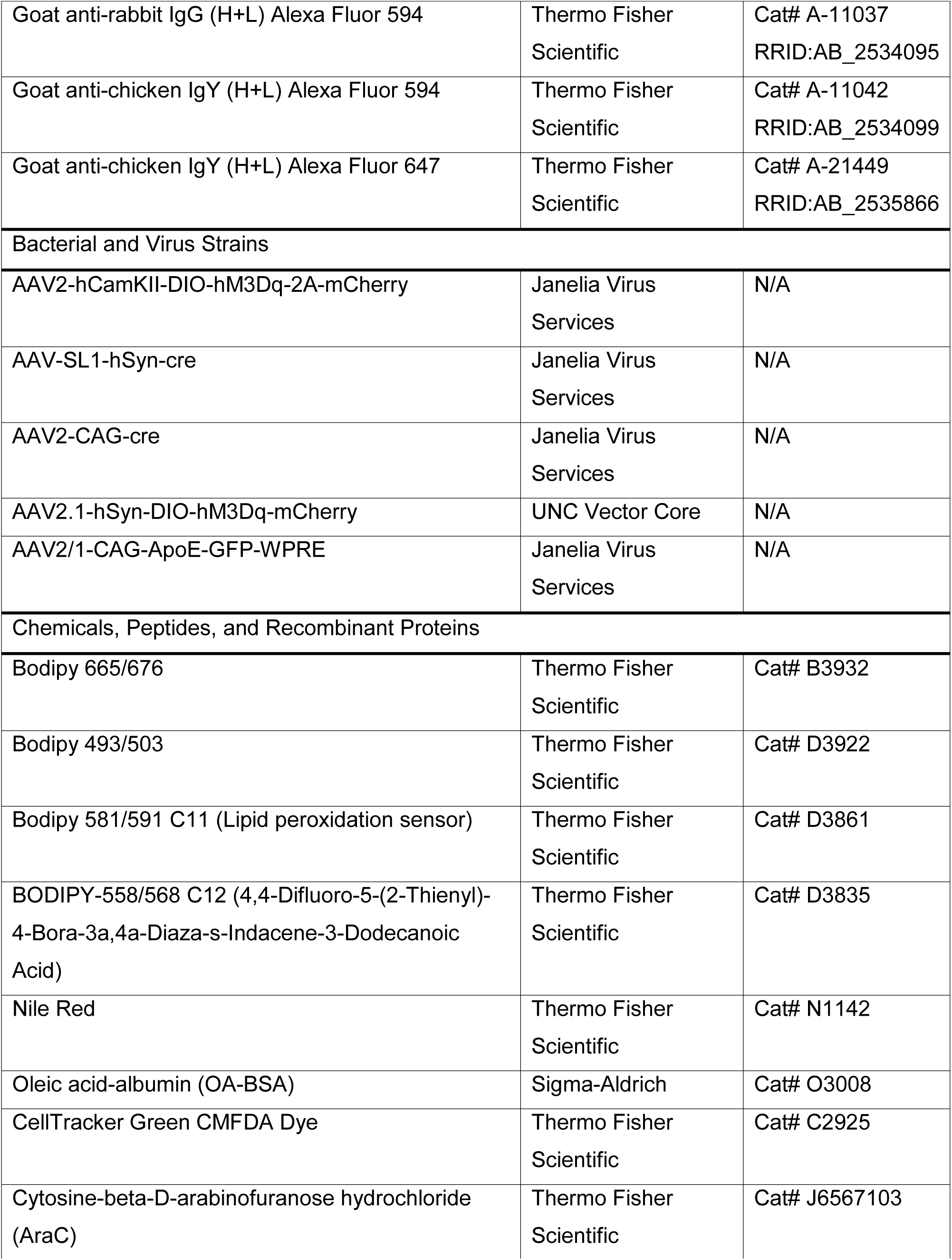
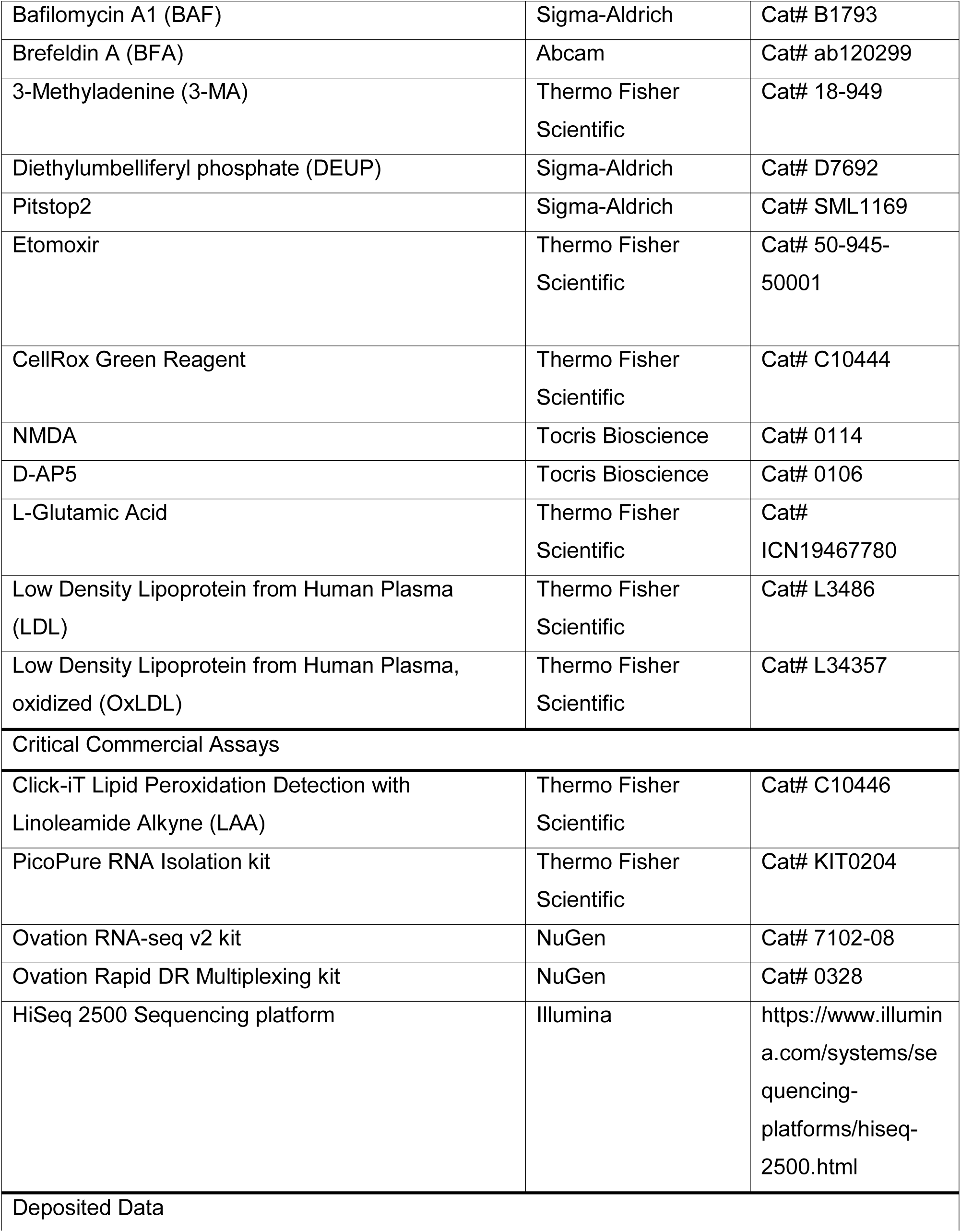
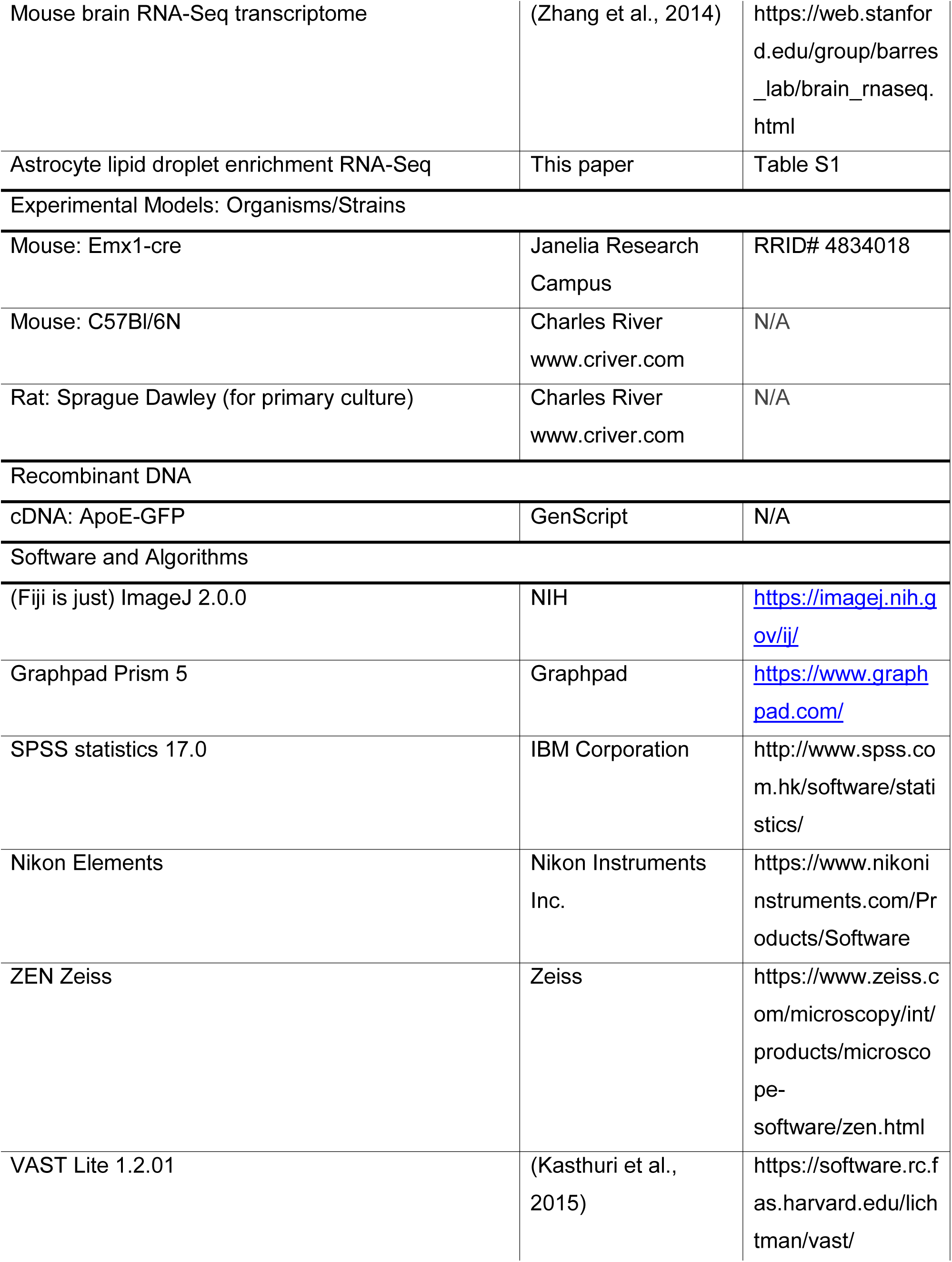
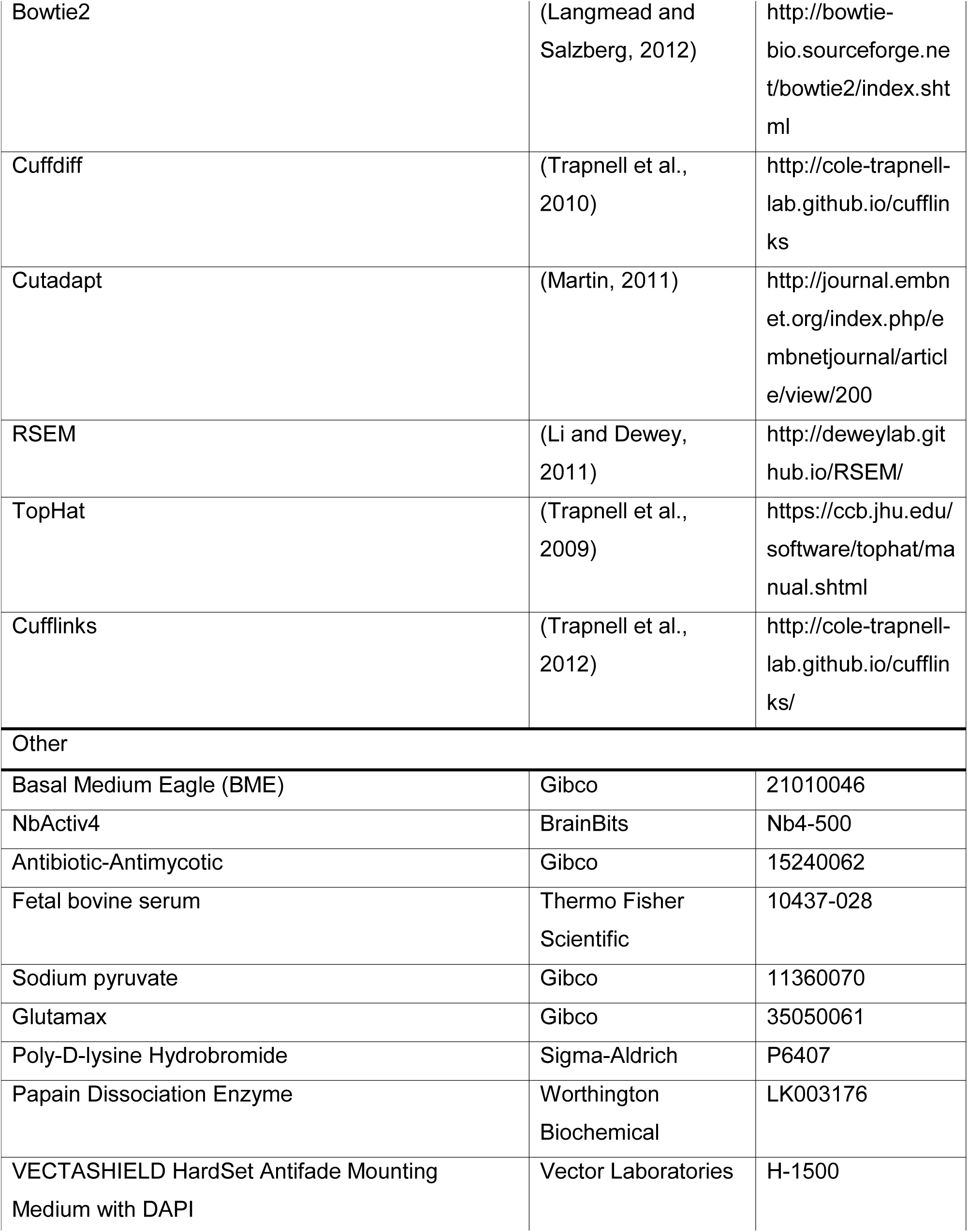
KEY RESOURCES TABLE

**KEY RESOURCES TABLE**
| REAGENT or RESOURCE | SOURCE | IDENTIFIER |
| --- | --- | --- |
| Antibodies |  |  |
| Rabbit polyclonal anti-NeuN | EMD Millipore | Cat# ABN78<br>RRID:AB_1080794<br>5 |
| Mouse monoclonal anti-Tuj1 | Biolegend | Cat# 801202<br>RRID:AB_1006340<br>8 |
| Chicken polyclonal anti-GFAP | Abcam | Cat# ab4674<br>RRID: AB_304558 |
| Rabbit polyclonal anti-Iba1 | Wako Chemicals | Cat# 019-19741<br>RRID: AB_839504 |
| Goat polyclonal anti-ApoE | Millipore Sigma | Cat# 178479<br>RRID:AB_1068296<br>5 |
| Rabbit polyclonal anti-LC3b | Abcam | Cat# ab48394<br>RRID:AB_881433 |
| Rabbit polyclonal anti-TOMM20 | Abcam | Cat# ab186735<br>RRID:AB_2716623 |
| Rabbit monoclonal anti-c-fos | Cell Signaling<br>Technology | Cat# 2250<br>RRID:AB_2247211 |
| Donkey anti-rabbit IgG (H+L) Alexa Fluor 488 | Thermo Fisher<br>Scientific | Cat# A-21206<br>RRID:AB_2535792 |
| Donkey anti-goat IgG (H+L) Alexa Fluor 488 | Thermo Fisher<br>Scientific | Cat# A-11055<br>RRID:AB_2534102 |
| Goat anti-mouse IgG (H+L) Alexa Fluor 488 | Thermo Fisher<br>Scientific | Cat# A-11001<br>RRID:AB_2534069 |
| Goat anti-chicken IgY (H+L) Alexa Fluor 488 | Thermo Fisher<br>Scientific | Cat# A-11039<br>RRID:AB_2534096 |
| Bovine anti-goat IgG (H+L) DyLight 549 | Jackson<br>ImmunoResearch | 805-505-180 |
| Goat anti-rabbit IgG (H+L) Alexa Fluor 594 | Thermo Fisher Scientific | Cat# A-11037<br>RRID:AB_2534095 |
| Goat anti-chicken IgY (H+L) Alexa Fluor 594 | Thermo Fisher Scientific | Cat# A-11042<br>RRID:AB_2534099 |
| Goat anti-chicken IgY (H+L) Alexa Fluor 647 | Thermo Fisher Scientific | Cat# A-21449<br>RRID:AB_2535866 |
| Bacterial and Virus Strains |  |  |
| AAV2-hCamKII-DIO-hM3Dq-2A-mCherry | Janelia Virus Services | N/A |
| AAV-SL1-hSyn-cre | Janelia Virus Services | N/A |
| AAV2-CAG-cre | Janelia Virus Services | N/A |
| AAV2.1-hSyn-DIO-hM3Dq-mCherry | UNC Vector Core | N/A |
| AAV2/1-CAG-ApoE-GFP-WPRE | Janelia Virus Services | N/A |
| Chemicals, Peptides, and Recombinant Proteins |  |  |
| Bodipy 665/676 | Thermo Fisher Scientific | Cat# B3932 |
| Bodipy 493/503 | Thermo Fisher Scientific | Cat# D3922 |
| Bodipy 581/591 C11 (Lipid peroxidation sensor) | Thermo Fisher Scientific | Cat# D3861 |
| BODIPY-558/568 C12 (4,4-Difluoro-5-(2-Thienyl)-4-Bora-3a,4a-Diaza-s-Indacene-3-Dodecanoic Acid) | Thermo Fisher Scientific | Cat# D3835 |
| Nile Red | Thermo Fisher Scientific | Cat# N1142 |
| Oleic acid-albumin (OA-BSA) | Sigma-Aldrich | Cat# O3008 |
| CellTracker Green CMFDA Dye | Thermo Fisher Scientific | Cat# C2925 |
| Cytosine-beta-D-arabinofuranose hydrochloride (AraC) | Thermo Fisher Scientific | Cat# J6567103 |
| Bafilomycin A1 (BAF) | Sigma-Aldrich | Cat# B1793 |
| Brefeldin A (BFA) | Abcam | Cat# ab120299 |
| 3-Methyladenine (3-MA) | Thermo Fisher Scientific | Cat# 18-949 |
| Diethylumbelliferyl phosphate (DEUP) | Sigma-Aldrich | Cat# D7692 |
| Pitstop2 | Sigma-Aldrich | Cat# SML1169 |
| Etomoxir | Thermo Fisher Scientific | Cat# 50-945-50001 |
| CellRox Green Reagent | Thermo Fisher Scientific | Cat# C10444 |
| NMDA | Tocris Bioscience | Cat# 0114 |
| D-AP5 | Tocris Bioscience | Cat# 0106 |
| L-Glutamic Acid | Thermo Fisher Scientific | Cat# ICN19467780 |
| Low Density Lipoprotein from Human Plasma (LDL) | Thermo Fisher Scientific | Cat# L3486 |
| Low Density Lipoprotein from Human Plasma, oxidized (OxLDL) | Thermo Fisher Scientific | Cat# L34357 |
| Critical Commercial Assays |  |  |
| Click-iT Lipid Peroxidation Detection with Linoleamide Alkyne (LAA) | Thermo Fisher Scientific | Cat# C10446 |
| PicoPure RNA Isolation kit | Thermo Fisher Scientific | Cat# KIT0204 |
| Ovation RNA-seq v2 kit | NuGen | Cat# 7102-08 |
| Ovation Rapid DR Multiplexing kit | NuGen | Cat# 0328 |
| HiSeq 2500 Sequencing platform | Illumina | https://www.illumina.com/systems/sequencing-platforms/hiseq-2500.html |
| Deposited Data |  |  |
| Mouse brain RNA-Seq transcriptome | (Zhang et al., 2014) | https://web.stanford.edu/group/barres_lab/brain_rnaseq.html |
| Astrocyte lipid droplet enrichment RNA-Seq | This paper | Table S1 |
| Experimental Models: Organisms/Strains |  |  |
| Mouse: Emx1-cre | Janelia Research Campus | RRID# 4834018 |
| Mouse: C57Bl/6N | Charles River<br>www.criver.com | N/A |
| Rat: Sprague Dawley (for primary culture) | Charles River<br>www.criver.com | N/A |
| Recombinant DNA |  |  |
| cDNA: ApoE-GFP | GenScript | N/A |
| Software and Algorithms |  |  |
| (Fiji is just) ImageJ 2.0.0 | NIH | https://imagej.nih.gov/ij/ |
| Graphpad Prism 5 | Graphpad | https://www.graphpad.com/ |
| SPSS statistics 17.0 | IBM Corporation | http://www.spss.com.hk/software/statistics/ |
| Nikon Elements | Nikon Instruments Inc. | https://www.nikoninstruments.com/Products/Software |
| ZEN Zeiss | Zeiss | https://www.zeiss.com/microscopy/int/products/microscope-software/zen.html |
| VAST Lite 1.2.01 | (Kasthuri et al., 2015) | https://software.rc.fas.harvard.edu/lichtman/vast/ |
| Bowtie2 | (Langmead and Salzberg, 2012) | http://bowtie-bio.sourceforge.net/bowtie2/index.shtml |
| Cuffdiff | (Trapnell et al., 2010) | http://cole-trapnell-lab.github.io/cufflinks |
| Cutadapt | (Martin, 2011) | http://journal.embn et.org/index.php/embnetjournal/article/view/200 |
| RSEM | (Li and Dewey, 2011) | http://deweylab.github.io/RSEM/ |
| TopHat | (Trapnell et al., 2009) | https://ccb.jhu.edu/software/tophat/manual.shtml |
| Cufflinks | (Trapnell et al., 2012) | http://cole-trapnell-lab.github.io/cufflinks/ |
| Other |  |  |
| Basal Medium Eagle (BME) | Gibco | 21010046 |
| NbActiv4 | BrainBits | Nb4-500 |
| Antibiotic-Antimycotic | Gibco | 15240062 |
| Fetal bovine serum | Thermo Fisher Scientific | 10437-028 |
| Sodium pyruvate | Gibco | 11360070 |
| Glutamax | Gibco | 35050061 |
| Poly-D-lysine Hydrobromide | Sigma-Aldrich | P6407 |
| Papain Dissociation Enzyme | Worthington Biochemical | LK003176 |
| VECTASHIELD HardSet Antifade Mounting Medium with DAPI | Vector Laboratories | H-1500 |

### Primary culture of hippocampal neurons and astrocytes

Primary hippocampal neurons were prepared as previously described (Beaudoin, III *et al.*, 2012). Briefly, the hippocampi were dissected from P0-P1 Sprague-Dawley rat pups and digested with papain (Worthington Biochemical). After digestion, the tissues were gently triturated and filtered with the cell strainer. Neurons were grown on poly-D-lysine coated glass coverslips in NbActiv4 medium containing antibiotic-antimycotic at 37 °C and used at DIV 5-7. Where specified 1 μm AraC was added to the media 2 days after plating and maintained for 4 days to prevent growth of astrocytes in neuronal cultures. Astrocyte cultures were grown in Basal Eagle Media containing 10% fetal bovine serum, 0.45% glucose, 1 mM sodium pyruvate, 2 mM glutamax supplement and antibiotic-antimycotic and used at DIV 3-5. For quantification, 5 images per coverslip were averaged.

### Cloning ApoE-GFP

ApoE-GFP cDNA was synthesized and was subcloned into the AAV-CAG-WPRE vector using BamHI and HindIII restriction sites:

GGATCCGCCACCATGAAGGTTCTGTGGGCTGCGTTGCTGGTCACATTCCTGGCAGGATGCCAGGCCAAGGTGGAGCAAGCGGTGGAGACAGAGCCGGAGCCCGAGCTGCGCCAGCAGACCGAGTGGCAGAGCGGCCAGCGCTGGGAACTGGCACTGGGTCGCTTTTGGGATTACCTGCGCTGGGTGCAGACACTGTCTGAGCAGGTGCAGGAGGAGCTGCTCAGCTCCCAGGTCACCCAGGAACTGAGGGCGCTGATGGACGAGACCATGAAGGAGTTGAAGGCCTACAAATCGGAACTGGAGGAACAACTGACCCCGGTGGCGGAGGAGACGCGGGCACGGCTGTCCAAGGAGCTGCAGGCGGCGCAGGCCCGGCTGGGCGCGGACATGGAGGACGTGTGCGGCCGCCTGGTGCAGTACCGCGGCGAGGTGCAGGCCATGCTCGGCCAGAGCACCGAGGAGCTGCGGGTGCGCCTCGCCTCCCACCTGCGCAAGCTGCGTAAGCGGCTCCTCCGCGATGCCGATGACCTGCAGAAGCGCCTGGCAGTGTACCAGGCCGGGGCCCGCGAGGGCGCCGAGCGCGGCCTCAGCGCCATCCGCGAGCGCCTGGGGCCCCTGGTGGAACAGGGCCGCGTGCGGGCCGCCACTGTGGGCTCCCTGGCCGGCCAGCCGCTACAGGAGCGGGCCCAGGCCTGGGGCGAGCGGCTGCGCGCGCGGATGGAGGAGATGGGCAGCCGGACCCGCGACCGCCTGGACGAGGTGAAGGAGCAGGTGGCGGAGGTGCGCGCCAAGCTGGAGGAGCAGGCCCAGCAGATACGCCTGCAGGCCGAGGCCTTCCAGGCCCGCCTCAAGAGCTGGTTCGAGCCCCTGGTGGAAGACATGCAGCGCCAGTGGGCCGGGCTGGTGGAGAAGGTGCAGGCTGCCGTGGGCACCAGCGCCGCCCCTGTGCCCAGCGACAATCACACCGGTATGGTGAGCAAGGGCGAGGAGCTGTTCACCGGGGTGGTGCCCATCCTGGTCGAGCTGGACGGCGACGTAAACGGCCACAAGTTCAGCGTGTCCGGCGAGGGCGAGGGCGATGCCACCTACGGCAAGCTGACCCTGAAGTTCATCTGCACCACCGGCAAGCTGCCCGTGCCCTGGCCCACCCTCGTGACCACCCTGACCTACGGCGTGCAGTGCTTCAGCCGCTACCCCGACCACATGAAGCAGCACGACTTCTTCAAGTCCGCCATGCCCGAAGGCTACGTCCAGGAGCGCACCATCTTCTTCAAGGACGACGGCAACTACAAGACCCGCGCCGAGGTGAAGTTCGAGGGCGACACCCTGGTGAACCGCATCGAGCTGAAGGGCATCGACTTCAAGGAGGACGGCAACATCCTGGGGCACAAGCTGGAGTACAACTACAACAGCCACAACGTCTATATCATGGCCGACAAGCAGAAGAACGGCATCAAGGTGAACTTCAAGATCCGCCACAACATCGAGGACGGCAGCGTGCAGCTCGCCGACCACTACCAGCAGAACACCCCCATCGGCGACGGCCCCGTGCTGCTGCCCGACAACCACTACCTGAGCACCCAGTCCG CCCTGAGCAAAGACCCCAACGAGAAGCGCGATCACATGGTCCTGCTGGAGTTCGTGACCGCCGCCGGGATCACTCTCGGCATGGACGAGCTGTACAAGTAAAAGCTT

### Confocal and widefield imaging

All fixed cells and tissue were mounted in vectashield HardSet Antifade Mounting Medium with DAPI. Imaging was performed using the following microscopes: (1) Nikon Eclipse TiE inverted microscope equipped with a 20x objective lens (Nikon, N.A. = 0.75) and sCMOS camera (Zyla 4.2, Andor) and elements software. (2) Nikon spinning disk equipped with a 60x oil objective lens (Nikon, N.A. = 1.4) and EMCCD iXon Ultra Camera (Andor) and elements software. (3) Keyence BZ-X700 inverted microscope equipped with a 20x objective lens (Nikon, N.A. = 0.75) and 3CCD camera. (4) 880 Laser Scanning Confocal Microscope (LSM) equipped with a plan-aprochromat 63x oil objective (Zeiss, NA=1.4) and 40x oil objective (Zeiss, NA=1.2) and ZEN software (Zeiss).

### Mitochondria fragmentation and surface area assay

Hippocampal neurons were treated with or without 500 μM NMDA in Nbactiv4 for 4 hours, fixed with 4% PFA, immunostained for TOMM20 to label mitochondria and imaged using the Zeiss 880 LSM. Images of Tomm20 were blindly scored based on mitochondrial fragmentation. Cells were scored as either “fragmented” or “non-fragmented”. The standard deviation was calculated based on the binomial distribution, sqrt(np(1-p)) where n is the number of cells and p the fraction of fragmented cells. For mitochondrial surface area, single confocal slices with the highest mitochondrial signal were used for quantification. Within ImageJ, a mask of the cell outline was made such that all pixels inside the neuron were equal to one and all pixels outside were null. Similarly, the mitochondria were thresholded such that all pixels inside the mitochondria were one and all pixels outside were null. The sum of all pixels was then measured for both images, essentially reporting the area occupied by either the total cell footprint or mitochondrial footprint. The ratio of the two is reported.

### Autophagy induction assay

Hippocampal neurons were treated with or without 500 μM NMDA in Nbactiv4 for 4 hours, fixed in ice-cold methanol, immunostained for LC3b to label autophagy and imaged using the Zeiss 880 LSM. Maximum intensity projections of three-dimensional image stacks of LC3b staining were first obtained. Then a series of image processing steps were performed to reduce background noise in order to automatically count LC3 positive puncta. Regions outside of the cell boundary were forced to null intensity to mediate false positive counts. Using a rolling ball radius of 5-20 pixels and sliding parabola the background of the image was then subtracted. Lastly, LC3 puncta were identified using the 3D Objects Counter (Bolte and Cordelieres, 2006) and the number of puncta per cell are reported.

### Lipid droplet assay

Hippocampal neurons or astrocytes washed in PBS and treated with or without the following drugs: 100 μM NMDA (500 μM NMDA if indicated on figure), 100 μM AP5,100 μM glutamate, 250 nM Bafilomycin A1 (BAF), 100 μM Diethylumbelliferyl phosphate (DEUP), 10 mM 3-Methyladenine (3MA), 300 nM etoxomir, 5 μg/mL Low Density Lipoprotein (LDL), or 5 μg/mL oxidized Low Density Lipoprotein (oxLDL) in artificial cerebral spinal fluid (125 mM NaCl, 5 mM KCl, 2 mM MgSO_4_, 24 mM NaHCO_3_, 1.25 NaH2PO_4_, 2 mM CaCl_2_ pH 7.4) containing 10 mM glucose unless otherwise indicated, for 4 hours at 37 °C. Cells were washed 3 times in PBS, fixed in 4% PFA and stained with 1 μg/mL Nile Red or 5 μg/mL BD 493/503 for 10 minutes at room temperature. Images for figures were taken using the Zeiss 880 LSM. Images for quantification were taken using the Nikon TiE widefield. Maximum intensity projections of three-dimensional image stacks of LD staining were first obtained. To remove background, a Gaussian blur was applied to a duplicate image and subtracted from the original image. The images were thresholded and particles were detected with a pixel size 2 to 100 and circularity from 0 to 1. For quantification, 10 images per coverslip were averaged.

### Fatty acid and ApoE transfer assays

Neurons were incubated with NbActiv4 containing 2 μM Bodipy 558/568 C_12_ (Red-C12) for 16 hours. Neurons were washed three times in warm PBS and incubated for an additional 1 hour in NbActiv4. Labelled neurons and unlabeled astrocytes on separate coverslips were washed twice with warm PBS and the coverslips were sandwiched together (facing each other) separated by paraffin wax and incubated in artificial cerebral spinal fluid (125 mM NaCl, 5 mM KCl, 2 mM MgSO_4_, 24 mM NaHCO_3_, 1.25 NaH2PO_4_, 2 mM CaCl_2_, 10 mM glucose pH 7.4) for 4 hours at 37 °C with or without 100 μM NMDA, 25 μM Pitstop2, 5 μg/mL Brefeldin A (BFA), 5 μg/mL Low Density Lipoprotein (LDL), or 5 μg/mL oxidized Low Density Lipoprotein (oxLDL). To test ApoE transfer, neurons DIV2 were transduced with AAV2/1-CAG-ApoE-GFP-WPRE (MOI = 1.5 × 105) and maintain for 10 days before using in transfer assay. To test the effects of neuronal activity, neurons DIV2 were transduced with AAV2-hSyn-DIO-hM3Dq-mCherry and AAV-SL1-CAG-Cre (MOI = 1.5 × 105) and maintain for 10 days before using in transfer assay with or without 1 μM clozapine-N-oxide for 4 hours. Cells were fixed and stained with Bodipy 493/503 to assess LDs in astrocytes and DAPI to assess cell death in neurons. Neurons expressing hM3Dq-mCherry were treated with or without 1 μM clozapine-N-oxide were fixed after 90 minutes and immunostained with anti-c-fos to confirm activation. Images for quantification were taken using the Nikon TiE widefield. Quantification of transfer was the same as LD assay.

### Fatty acid pulse-chase assays

Astrocytes were incubated in complete media containing 2 μM Bodipy 558/568 C_12_ (Red-C12) for 16 hours. Astrocytes were washed three times in warm PBS and incubated for an additional 1 hour in complete media and chased in aCSF with or without 100 μM NMDA. Astrocytes were fixed and mitochondria were immunolabelled with anti-Tomm20 to visualize mitochondria. Imaging was performed on the Zeiss 880 LSM. Quantification of transfer was the same as LD assay.

### Lipoprotein particle depletion assay

Neurons were incubated with NbActiv4 containing 2 μM Bodipy 558/568 C_12_ (Red-C12) for 16 hours. Neurons were washed three times in warm PBS, incubated for an additional 1 hour in NbActiv4 followed by 4 hours in aCSF at 37 °C. Neuron conditioned aCSF was collected and passed through a 0.2 μm filter, or centrifuged for 4000 × g for 10 min, then 10,000 × g for 30 min then 250,000 × g for 2.5 hr at 4°C. Astrocytes were washed 3 times in PBS and incubated with deleted neuron-conditioned aCSF for 4 hours in aCSF at 37 °C, fixed, mounted and imaged on the Zeiss 880 LSM. Quantification of transfer was the same as LD assay. Neuron-conditioned aCSF or aCSF containing 10 μM oleic-acid bound to albumin was centrifuged and analyzed by western blot. Samples were centrifuged for 4000 × g for 10 min and 10 μl was analyzed as starting material. The supernatant was centrifuged at 10,000 × g for 30 min (low g) followed by 250,000 × g for 2.5 hr (high g) at 4°C and10 μl of supernatants and pellets resuspended in 10 μl aCSF were analyzed by SDS-PAGE and Western blotting. Oleic-acid bound to albumin was analyzed by Coomassie blue stain.

### Lipid peroxidation and ROS Assays

For Click-iT lipid peroxidation assay, neurons were treated for 4 hours in NbActiv4 with or without 500 μM NMDA with 50 mM linoleamide alkyne added for the last 2 hours. Cells were washed three times in PBS, fixed in 4 % paraformaldehyde for 10 min and blocked in 1% BSA for 30 minutes. Cells were incubated in the Click-iT reaction cocktail for 30 minutes at room temperature, washed twice with 2% BSA, twice with PBS, mounted and imaged using the Zeiss 880 LSM. Mean intensities and integrated density for each thresholded cell were measured using ImageJ. Quantification of transfer was the same as LD assay. For quantification, 8 images per coverslip were averaged. For CellRox assay, astrocytes were treated for 4 hours in aCSF with or without 100 μM NMDA with 5 μM CellROX Green added for the last 30 min. Cells were washed three times with PBS and fixed with 3 % PFA for 10 minutes and mounted. Imaging was performed on a Nikon Eclipse TiE Inverted microscope equipped with a 60X Oil-immersion objective lens (Nikon, N.A. = 1.4) and EMCCD camera (iXon Ultra 897, Andor). Images were shade corrected and mean intensities and integrated density for each thresholded nuclei were measured using ImageJ. For quantification, 10 images per coverslip were averaged For Bodipy-C11 lipid peroxidation assay, astrocytes were incubated in HBSS with or without 100 μM NMDA for 4 hr with 2 μM C11-BODIPY added for the last 30 min. Cells were then wash with HBSS and imaged live in HBSS with or without NMDA using the Nikon spinning disk confocal microscope. Z-stacks were taken at both green and red channels. Relative lipid peroxidation was indicated by the ratio of green fluorescence intensity over red fluorescence intensity using imageJ.

### Mouse brain sample preparation for FIB-SEM

2-month old male C57/BL6 mouse was deeply anesthetized and transcardially perfused with 30 ml of 3% PFA (60 mM NaCl, 130 mM glycerol, 10 mM sodium phosphate buffer). The brain was carefully dissected out from the skull and postfixed with 50 ml of 3% PFA (30 mM of NaCl, 70 mM glycerol, 30 mM PIPES buffer, 10 mM betaine, 2 mM CaCl_2_, 2 mM MgSO_4_) under room temperature for 2 hours. The brain sample was then rinsed in a 400 mOsM buffer (65 mM NaCl, 100 mM glycerol, 30 mM PIPES buffer, 10 mM betaine, 2 mM CaCl_2_, and 2 mM MgSO_4_) for half an hour, followed by vibratome sectioning (coronal sections, 100 μm thickness) using a Leica VT1000S vibratome in the same buffer. 100 μm sections were then fixed in 1% PFA, 2% glutaraldehyde solution (30 mM NaCl, 70 mM glycerol, 30 mM PIPES buffer, 10 mM betaine, 2 mM CaCl_2_, 2 mM MgSO_4_, 75 mM sucrose) overnight at 4 degrees. Sections were then washed using the 400 mOsM rinsing buffer (see above). Round samples of the hippocampus were created from the 100 μm coronal sections using a 2 mm biopsy punch (Miltex). The 2 mm samples were dipped in hexadecene, placed in 100 μm aluminum carrier, covered with a flat carrier and high-pressure frozen using a Wohlwend compact 01 High pressure freezer. Samples were then freeze-substituted in 0.5% osmium tetroxide, 20 mM 3-Amino-1,2,4-triazole, 0.1% uranyl acetate, 4% water in acetone, using a Leica AFS2 system. Specimens were further dehydrated in 100% acetone and embedded in Durcupan resin.

### FIB-SEM imaging of mouse brain sample

Durcupan embedded mouse hippocampus CA1 sample was first mounted on a Cu stud, then imaged by a customized Zeiss Merlin FIB-SEM system previously described (Xu, C. S. et. al. Enhanced FIB-SEM Systems for Large-Volume 3D Imaging, eLife 2017 May; doi: 10.7554/eLife.25916). The block face was imaged by a 2 nA electron beam with 1.2 keV landing energy at 2 MHz. The x-y pixel resolution was set at 8 nm. A subsequently applied focused Ga+ beam of 15 nA at 30 keV strafed across the top surface and ablated away 4 nm of the surface. The newly exposed surface was then imaged again. The ablation – imaging cycle continued about once every minute for one week. The sequence of acquired images formed a raw imaged volume, followed by post processing of image registration and alignment using a Scale Invariant Feature Transform (SIFT) based algorithm. The aligned stack was binned by a factor of 2 along z to form a final isotropic volume of 60 × 80 × 45 μm^3^ with 8 × 8 × 8 nm^3^ voxels, which can be viewed at any arbitrary orientations.

### Cell Sorting and RNA-seq library preparation

Primary hippocampal astrocytes (DIV 4) were incubated for 10 minutes in 1 ng/mL Nile Red in PBS at 37 °C, trypsinized to detach from the cultured cells, spun at 250 × G for 5 minutes, re-suspended in PBS and passed through a cell strainer to yield single cells. Cells were sorted for on the BD FACS ARIA II using the 488nm laser to separate LD positive and negative cells. Unstained cells were used as a negative control to establish the gates and exclude dead cells and doublets. Sorted astrocytes were lysed in PicoPure extraction buffer. RNA extraction and RNA library preparation was performed as described in (Cembrowski et al., 2016). Briefly, RNA was extracted using PicoPure RNA Isolation kit, cDNA was amplified using Ovation RNA-seq v2 kit and the Ovation Rapid DR Multiplexing kit was used to make the sequencing library. Libraries were sequenced on the HiSeq 2500 platform with single-end 100 bp reads. Reads were first trimmed using cutadapt (Martin, 2011), then aligned to the rn6 transcriptome (GCF_000001895.5) using RSEM (Li and Dewey, 2011). Gene expression estimates were reported as Fragments Per Kilobase of transcript per Million mapped reads (FPKM).

### Gene expression level testing and gene ontology analysis

We mapped sequencing data back to the rat reference genome (Rt6) by Bowtie2 (Langmead and Salzberg, 2012). Transcript isoforms were reconstructed from PE reads using TopHat (Trapnell et al., 2009) and abundances estimated using Cufflinks (Trapnell et al., 2012). Read counts were tallied for each Ensembl annotated protein-coding gene incremented by 1 and differential expression tested using Cuffdiff using all qualified samples. Gene Ontology analysis was performed separately on up-regulated and down-regulated genes using DAVID Bioinformatics Functional Annotation Tool (Huang et al., 2007b; Huang et al., 2007a). Cell-type specific mouse mRNA seq data for astrocyte, neuron, oligodendrocyte progenitor cells, newly-formed oligodendrocytes, myelinating oligodendrocytes, microglia and endothelial cells were obtained from (Zhang et al., 2014). Cross-species relative gene expression levels were calculated by direct normalization of FPKM of homologous genes.

### Pial strip devascularization and chemogenetic stimulations

All procedures were conducted in accordance with protocols approved by the Janelia Institutional Animal Care. Pial strip devascularization was performed as previously described (Farr and Whishaw, 2002; Alaverdashvili et al., 2008; Karl et al., 2010). Adult C57BL6 mice were anaesthetized with isoflurane and buprenorphine (0.1 mg/kg, SC) was administered as an analgesic. A craniometry was performed over the motor cortex. The coordinates of the square incised measured from Bregma; (a) A +1.0 mm, L 1.0 mm (b) A 1.0 mm, L 2.0 mm (c) A 2.0 mm, L 2.0 mm (d) A 2.0 mm, L 1.0 mm. The dura mater was cut and peeled away using the sharp edge of a sterile hypodermic needle and fine forceps. Sterile saline-soaked absorption spears were used to wipe away the pia and superficial vasculature, the area was flushed with ample sterile saline and gelfoam was applied to the stroke region which aids in coagulation. A thin layer of artificial dura and bone wax was applied to protect the exposed tissue and the wound was closed with stitches and vetbond. Marcaine was applied to the incision site and ketoprofen (5 mg/kg) was administered at the end of the surgery and once per day for 2 days after surgery. For in vivo chemogenetic stimulation, mice were anesthetized with isoflurane and a small craniotomy (0.1 mm × 0.1 mm) was performed over the motor cortex for in the insertion of the injection needle. Emx1-cre mice were injected with 200 nL of AAV2.1-hSyn-DIO-hM3Dq-mCherry and C57BL6 mice were co-injected with AAV2-hCamKII-DIO-hM3Dq-2A-mCherry or AAV2.1-hSyn-DIO-hM3Dq-mCherry plus 200 nL AAV-SL1-hSyn-cre into the motor cortex (coordinates A/P: 1.0 mm, M/L: 1.5 mm, D/V: 0.5 mm). All virus titers were 10^13^ GC/mL. Marcaine was applied to the incision site and ketoprofen (1 mg/kg) was administered at the end of the surgery and once per day for 2 days after surgery. 4 weeks post-injection, mice were injected with clozapine-N-oxide (5 mg/kg) or saline intraperitoneally, twice a day for three days. For quantification, 3-5 images per hemisphere or treatment were averaged

### Histology

Mice were deeply anesthetized with isoflurane and intracardially perfused with 4% paraformaldehyde in phosphate-buffered saline. The brains were removed and post-fixed in 4% PFA overnight at 4 °C. Serial 70 μm sections spanning the lesion or injection site were obtained on a vibratome. Slices were incubated at room temperature for 1 hr in blocking buffer (PBS, 2% BSA, 0.2% Triton-X-100), overnight in primary antibody in blocking buffer and 4 hrs in secondary antibody plus Bodipy-493/503 in blocking buffer. Between antibodies, slices were washed 3 times with PBS containing 0.2% TritonX-100. Imaging was performed using the Nikon TiE and Keyence for whole brain imaging and stitching. Cell type and subcellular imaging was performed using the Zeiss 880 LSM.

### Statistical analysis

Statistical analysis was performed using Graphpad Prism 5 and SPSS Statistics 17.0. Statistical significance of parametric datasets was determined by t-test for comparison of two groups or by ANOVA with post-hoc tukey test for multiple comparisons. Where control groups were normalized to 1, non-parametic Mann-Whitney U test for comparison of two groups or Kruskal-Wallis test and post-hoc Chi-square test for multiple comparisons

## Author Contributions

MSI, JLS and ZL designed experiments. MSI, CLC, AVW and HL performed and analyzed *in vitro* experiments. JJ and MSI performed and analyzed *in vivo* chemogenetic and pial lesion experiments. MSI and ZL analyzed RNAseq data. SHS and HAP prepared mouse brain sample for FIB-SEM. CSX, SP and HFH generated and processed the FIB-SEM dataset. SHS, MSI and HAP analyzed the FIB-SEM dataset. MSI, JLS and ZL wrote the manuscript.

## Acknowledgments

We thank Salvatore DiLisio, Anne Kuszpit and Colin Morrow for assistance with virus injections and animal care; Deepika Walpita for assistance with cell sorting and primary cell cultures; Andrew Lemire and Kshama Aswath for help with RNAseq data generation; Melissa Ramirez and Jordon Towne for cloning the ApoE-GFP construct; Janelia Virus Services for producing viruses; Melanie Radcliff for administrative assistance; Jeremy Cohen, Carolyn Ott, Sarah Cohen and Andrea Marat for helpful discussion and comments on the manuscript. The work was supported by the Howard Hughes Medical Institute.

## Reference List

Adamsky, A., Kol, A., Kreisel, T., Doron, A., Ozeri-Engelhard, N., Melcer, T., Refaeli, R., Horn, H., Regev, L., Groysman, M., London, M., and Goshen, I. (2018). Astrocytic Activation Generates De Novo Neuronal Potentiation and Memory Enhancement. Cell 174, 59–71.

Aguado, F., Espinosa-Parrilla, J.F., Carmona, M.A., and Soriano, E. (2002). Neuronal activity regulates correlated network properties of spontaneous calcium transients in astrocytes in situ. J. Neurosci. 22, 9430–9444.

Alaverdashvili, M., Moon, S.K., Beckman, C.D., Virag, A., and Whishaw, I.Q. (2008). Acute but not chronic differences in skilled reaching for food following motor cortex devascularization vs. photothrombotic stroke in the rat. Neuroscience 157, 297–308.

Alexander, G.M., Rogan, S.C., Abbas, A.I., Armbruster, B.N., Pei, Y., Allen, J.A., Nonneman, R.J., Hartmann, J., Moy, S.S., Nicolelis, M.A., McNamara, J.O., and Roth, B.L. (2009). Remote control of neuronal activity in transgenic mice expressing evolved G protein-coupled receptors. Neuron 63, 27–39.

Anderson, C.M. and Swanson, R.A. (2000). Astrocyte glutamate transport: review of properties, regulation, and physiological functions. Glia 32, 1–14.

Aoki, K., Uchihara, T., Sanjo, N., Nakamura, A., Ikeda, K., Tsuchiya, K., and Wakayama, Y. (2003). Increased expression of neuronal apolipoprotein E in human brain with cerebral infarction. Stroke 34, 875–880.

Bailey, A.P., Koster, G., Guillermier, C., Hirst, E.M., MacRae, J.I., Lechene, C.P., Postle, A.D., and Gould, A.P. (2015). Antioxidant Role for Lipid Droplets in a Stem Cell Niche of Drosophila. Cell 163, 340–353.

Beaudoin, G.M., III, Lee, S.H., Singh, D., Yuan, Y., Ng, Y.G., Reichardt, L.F., and Arikkath, J. (2012). Culturing pyramidal neurons from the early postnatal mouse hippocampus and cortex. Nat. Protoc. 7, 1741–1754.

Belanger, M., Allaman, I., and Magistretti, P.J. (2011). Brain energy metabolism: focus on astrocyte-neuron metabolic cooperation. Cell Metab 14, 724–738.

Belanger, M. and Magistretti, P.J. (2009). The role of astroglia in neuroprotection. Dialogues. Clin. Neurosci. 11, 281–295.

Ben, H.L., Carrillo-de Sauvage, M.A., Ceyzeriat, K., and Escartin, C. (2015). Elusive roles for reactive astrocytes in neurodegenerative diseases. Front Cell Neurosci. 9, 278.

Bolte, S. and Cordelieres, F.P. (2006). A guided tour into subcellular colocalization analysis in light microscopy. J. Microsc. 224, 213–232.

Bowser, D.N. and Khakh, B.S. (2004). ATP excites interneurons and astrocytes to increase synaptic inhibition in neuronal networks. J. Neurosci. 24, 8606–8620.

Buttini, M., Masliah, E., Yu, G.Q., Palop, J.J., Chang, S., Bernardo, A., Lin, C., Wyss-Coray, T., Huang, Y., and Mucke, L. (2010). Cellular source of apolipoprotein E4 determines neuronal susceptibility to excitotoxic injury in transgenic mice. Am. J. Pathol. 177, 563–569.

Cembrowski, M.S., Wang, L., Sugino, K., Shields, B.C., and Spruston, N. (2016). Hipposeq: a comprehensive RNA-seq database of gene expression in hippocampal principal neurons. Elife. 5, e14997.

Davies, J.P., Chen, F.W., and Ioannou, Y.A. (2000). Transmembrane molecular pump activity of Niemann-Pick C1 protein. Science 290, 2295–2298.

Dekroon, R.M. and Armati, P.J. (2001). Synthesis and processing of apolipoprotein E in human brain cultures. Glia 33, 298–305.

DeMattos, R.B., Curtiss, L.K., and Williams, D.L. (1998). A minimally lipidated form of cell-derived apolipoprotein E exhibits isoform-specific stimulation of neurite outgrowth in the absence of exogenous lipids or lipoproteins. J. Biol. Chem. 273, 4206–4212.

Dzamba, D., Honsa, P., and Anderova, M. (2013). NMDA Receptors in Glial Cells: Pending Questions. Curr. Neuropharmacol. 11, 250–262.

Farr, T.D. and Whishaw, I.Q. (2002). Quantitative and qualitative impairments in skilled reaching in the mouse (Mus musculus) after a focal motor cortex stroke. Stroke 33, 1869–1875.

Greenspan, P., Mayer, E.P., and Fowler, S.D. (1985). Nile red: a selective fluorescent stain for intracellular lipid droplets. J. Cell Biol. 100, 965–973.

Guo, Y., Cordes, K.R., Farese, R.V., Jr., and Walther, T.C. (2009). Lipid droplets at a glance. J. Cell Sci. 122, 749–752.

Haba, K., Ogawa, N., Mizukawa, K., and Mori, A. (1991). Time course of changes in lipid peroxidation, pre- and postsynaptic cholinergic indices, NMDA receptor binding and neuronal death in the gerbil hippocampus following transient ischemia. Brain Res. 540, 116–122.

Huang, D.W., Sherman, B.T., Tan, Q., Collins, J.R., Alvord, W.G., Roayaei, J., Stephens, R., Baseler, M.W., Lane, H.C., and Lempicki, R.A. (2007a). The DAVID Gene Functional Classification Tool: a novel biological module-centric algorithm to functionally analyze large gene lists. Genome Biol. 8, R183.

Huang, D.W., Sherman, B.T., Tan, Q., Kir, J., Liu, D., Bryant, D., Guo, Y., Stephens, R., Baseler, M.W., Lane, H.C., and Lempicki, R.A. (2007b). DAVID Bioinformatics Resources: expanded annotation database and novel algorithms to better extract biology from large gene lists. Nucleic Acids Res. 35, W169–W175.

Karl, J.M., Alaverdashvili, M., Cross, A.R., and Whishaw, I.Q. (2010). Thinning, movement, and volume loss of residual cortical tissue occurs after stroke in the adult rat as identified by histological and magnetic resonance imaging analysis. Neuroscience 170, 123–137.

Kasthuri, N., Hayworth, K.J., Berger, D.R., Schalek, R.L., Conchello, J.A., Knowles-Barley, S., Lee, D., Vazquez-Reina, A., Kaynig, V., Jones, T.R., Roberts, M., Morgan, J.L., Tapia, J.C., Seung, H.S., Roncal, W.G., Vogelstein, J.T., Burns, R., Sussman, D.L., Priebe, C.E., Pfister, H., and Lichtman, J.W. (2015). Saturated Reconstruction of a Volume of Neocortex. Cell 162, 648–661.

Kawamura, M., Gachet, C., Inoue, K., and Kato, F. (2004). Direct excitation of inhibitory interneurons by extracellular ATP mediated by P2Y1 receptors in the hippocampal slice. J. Neurosci. 24, 10835–10845.

Kim, W.S., Rahmanto, A.S., Kamili, A., Rye, K.A., Guillemin, G.J., Gelissen, I.C., Jessup, W., Hill, A.F., and Garner, B. (2007). Role of ABCG1 and ABCA1 in regulation of neuronal cholesterol efflux to apolipoprotein E discs and suppression of amyloid-beta peptide generation. J. Biol. Chem. 282, 2851–2861.

Klausner, R.D., Donaldson, J.G., and Lippincott-Schwartz, J. (1992). Brefeldin A: insights into the control of membrane traffic and organelle structure. J. Cell Biol. 116, 1071–1080.

Lai, T.W., Zhang, S., and Wang, Y.T. (2014). Excitotoxicity and stroke: Identifying novel targets for neuroprotection. Progress in Neurobiology 115, 157–188.

Langmead, B. and Salzberg, S.L. (2012). Fast gapped-read alignment with Bowtie 2. Nat. Methods 9, 357–359.

Li, B. and Dewey, C.N. (2011). RSEM: accurate transcript quantification from RNA-Seq data with or without a reference genome. BMC. Bioinformatics. 12, 323.

Lippincott-Schwartz, J., Yuan, L., Tipper, C., Amherdt, M., Orci, L., and Klausner, R.D. (1991). Brefeldin A’s effects on endosomes, lysosomes, and the TGN suggest a general mechanism for regulating organelle structure and membrane traffic. Cell 67, 601–616.

Liu, L., MacKenzie, K.R., Putluri, N., Maletic-Savatic, M., and Bellen, H.J. (2017). The Glia-Neuron Lactate Shuttle and Elevated ROS Promote Lipid Synthesis in Neurons and Lipid Droplet Accumulation in Glia via APOE/D. Cell Metab 26, 719–737.

Liu, L., Zhang, K., Sandoval, H., Yamamoto, S., Jaiswal, M., Sanz, E., Li, Z., Hui, J., Graham, B.H., Quintana, A., and Bellen, H.J. (2015). Glial lipid droplets and ROS induced by mitochondrial defects promote neurodegeneration. Cell 160, 177–190.

Mahley, R.W. (2016). Central Nervous System Lipoproteins: ApoE and Regulation of Cholesterol Metabolism. Arterioscler. Thromb. Vasc. Biol. 36, 1305–1315.

Martin, M. (2011). Cutadapt Removes Adapter Sequences From High-Throughput Sequencing Reads. EMBnet. journal 17, 10–12.

Nguyen, D., Alavi, M.V., Kim, K.Y., Kang, T., Scott, R.T., Noh, Y.H., Lindsey, J.D., Wissinger, B., Ellisman, M.H., Weinreb, R.N., Perkins, G.A., and Ju, W.K. (2011). A new vicious cycle involving glutamate excitotoxicity, oxidative stress and mitochondrial dynamics. Cell Death. Dis. 2, e240.

Nguyen, T.B., Louie, S.M., Daniele, J.R., Tran, Q., Dillin, A., Zoncu, R., Nomura, D.K., and Olzmann, J.A. (2017). DGAT1-Dependent Lipid Droplet Biogenesis Protects Mitochondrial Function during Starvation-Induced Autophagy. Dev. Cell 42, 9–21.

Parihar, M.S. and Hemnani, T. (2004). Experimental excitotoxicity provokes oxidative damage in mice brain and attenuation by extract of Asparagus racemosus. J. Neural Transm. (Vienna.) 111, 1–12.

Parthasarathy, S., Raghavamenon, A., Garelnabi, M.O., and Santanam, N. (2010). Oxidized low-density lipoprotein. Methods Mol. Biol. 610, 403–417.

Rambold, A.S., Cohen, S., and Lippincott-Schwartz, J. (2015). Fatty acid trafficking in starved cells: regulation by lipid droplet lipolysis, autophagy, and mitochondrial fusion dynamics. Dev. Cell 32, 678–692.

Redmann, M., Benavides, G.A., Berryhill, T.F., Wani, W.Y., Ouyang, X., Johnson, M.S., Ravi, S., Barnes, S., Darley-Usmar, V.M., and Zhang, J. (2017). Inhibition of autophagy with bafilomycin and chloroquine decreases mitochondrial quality and bioenergetic function in primary neurons. Redox. Biol. 11, 73–81.

Ren, Z., Iliff, J.J., Yang, L., Yang, J., Chen, X., Chen, M.J., Giese, R.N., Wang, B., Shi, X., and Nedergaard, M. (2013). ‘Hit & Run’ model of closed-skull traumatic brain injury (TBI) reveals complex patterns of post-traumatic AQP4 dysregulation. J. Cereb. Blood Flow Metab 33, 834–845.

Reynolds, I.J. and Hastings, T.G. (1995). Glutamate induces the production of reactive oxygen species in cultured forebrain neurons following NMDA receptor activation. J. Neurosci. 15, 3318–3327.

Rong, X., Wang, B., Dunham, M.M., Hedde, P.N., Wong, J.S., Gratton, E., Young, S.G., Ford, D.A., and Tontonoz, P. (2015). Lpcat3-dependent production of arachidonoyl phospholipids is a key determinant of triglyceride secretion. Elife. 4.

Schonfeld, P. and Reiser, G. (2013). Why does brain metabolism not favor burning of fatty acids to provide energy? Reflections on disadvantages of the use of free fatty acids as fuel for brain. J. Cereb. Blood Flow Metab 33, 1493–1499.

Schonfeld, P. and Reiser, G. (2017). Brain energy metabolism spurns fatty acids as fuel due to their inherent mitotoxicity and potential capacity to unleash neurodegeneration. Neurochem. Int. 109, 68–77.

Sultana, R., Perluigi, M., and Allan, B.D. (2013). Lipid peroxidation triggers neurodegeneration: a redox proteomics view into the Alzheimer disease brain. Free Radic. Biol. Med. 62, 157–169.

Trapnell, C., Pachter, L., and Salzberg, S.L. (2009). TopHat: discovering splice junctions with RNA-Seq. Bioinformatics. 25, 1105–1111.

Trapnell, C., Roberts, A., Goff, L., Pertea, G., Kim, D., Kelley, D.R., Pimentel, H., Salzberg, S.L., Rinn, J.L., and Pachter, L. (2012). Differential gene and transcript expression analysis of RNA-seq experiments with TopHat and Cufflinks. Nat. Protoc. 7, 562–578.

Trapnell, C., Williams, B.A., Pertea, G., Mortazavi, A., Kwan, G., van Baren, M.J., Salzberg, S.L., Wold, B.J., and Pachter, L. (2010). Transcript assembly and quantification by RNA-Seq reveals unannotated transcripts and isoform switching during cell differentiation. Nat. Biotechnol. 28, 511–515.

Unger, R.H., Clark, G.O., Scherer, P.E., and Orci, L. (2010). Lipid homeostasis, lipotoxicity and the metabolic syndrome. Biochim. Biophys. Acta 1801, 209–214.

Wang, H., Wei, E., Quiroga, A.D., Sun, X., Touret, N., and Lehner, R. (2010). Altered lipid droplet dynamics in hepatocytes lacking triacylglycerol hydrolase expression. Mol. Biol. Cell 21, 1991–2000.

Xu, C.S., Hayworth, K.J., Lu, Z., Grob, P., Hassan, A.M., Garcia-Cerdan, J.G., Niyogi, K.K., Nogales, E., Weinberg, R.J., and Hess, H.F. (2017). Enhanced FIB-SEM systems for large-volume 3D imaging. Elife. 6.

Xu, Q., Bernardo, A., Walker, D., Kanegawa, T., Mahley, R.W., and Huang, Y. (2006). Profile and regulation of apolipoprotein E (ApoE) expression in the CNS in mice with targeting of green fluorescent protein gene to the ApoE locus. J. Neurosci. 26, 4985–4994.

Yellen, G. (2018). Fueling thought: Management of glycolysis and oxidative phosphorylation in neuronal metabolism. J. Cell Biol. 217, 2235–2246.

Zeng, Y., Tao, N., Chung, K.N., Heuser, J.E., and Lublin, D.M. (2003). Endocytosis of oxidized low density lipoprotein through scavenger receptor CD36 utilizes a lipid raft pathway that does not require caveolin-1. J. Biol. Chem. 278, 45931–45936.

Zhang, J.M., Wang, H.K., Ye, C.Q., Ge, W., Chen, Y., Jiang, Z.L., Wu, C.P., Poo, M.M., and Duan, S. (2003). ATP released by astrocytes mediates glutamatergic activity-dependent heterosynaptic suppression. Neuron 40, 971–982.

Zhang, L., Song, J., Cavigiolio, G., Ishida, B.Y., Zhang, S., Kane, J.P., Weisgraber, K.H., Oda, M.N., Rye, K.A., Pownall, H.J., and Ren, G. (2011). Morphology and structure of lipoproteins revealed by an optimized negative-staining protocol of electron microscopy. J. Lipid Res. 52, 175–184.

Zhang, Y., Chen, K., Sloan, S.A., Bennett, M.L., Scholze, A.R., O’Keeffe, S., Phatnani, H.P., Guarnieri, P., Caneda, C., Ruderisch, N., Deng, S., Liddelow, S.A., Zhang, C., Daneman, R., Maniatis, T., Barres, B.A., and Wu, J.Q. (2014). An RNA-sequencing transcriptome and splicing database of glia, neurons, and vascular cells of the cerebral cortex. J. Neurosci. 34, 11929–11947.

